# Adaptive Partial Conjunction Hypothesis: Identifying Pleiotropy Across Heterogeneous Effect Units

**DOI:** 10.1101/2025.11.25.690340

**Authors:** Yuxin Li, Zicheng Lu, Xiaolei Lin

## Abstract

Genetic effects are increasingly measured across multiple traits, tissues and cell types. Variants can influence several traits through shared pathways and regulate gene expression across several tissues. Resolving these pleiotropic patterns can reveal shared genetic architecture and disease mechanisms. Addressing this calls for the partial conjunction hypothesis (PCH), namely that every feature in a subset is affected. Existing PCH methods, however, rely on simple marginal effect-size models and pool effect sizes within each feature, which is insufficient when effects are heterogeneous. We introduce APCH, an empirical Bayes framework for subset-level inference. APCH models each feature’s effect-size distribution with a flexible adaptive shrinkage prior and learns how effect sizes are shared among jointly affected features. We define subset-level posterior quantities, notably a local false sign rate that is more robust and interpretable. In simulations spanning direct pleiotropy, structural-equation effects and genetic correlation, APCH remained calibrated and powerful. Across five type 2 diabetes (T2D)-related traits, APCH identified 12 loci jointly associated with all five traits, whereas only one variant reached genome-wide significance in all five. Across six TWAS of T2D in different tissues, APCH recovered gene–tissue associations missed by single-tissue analyses.

## 1 Introduction

Pleiotropy, a central concept in genetics research, refers to the phenomenon where a single gene or genetic variant is associated with multiple phenotypes [1, 2]. For example, researchers are interested in testing whether a single locus is associated with multiple diseases to discover their shared genetic etiology [3–5]. Moreover, they investigate whether a single variant drives signals for both a clinical and a molecular phenotype, aiming to link statistical findings to biological mechanisms [6, 7]. These problems share a common goal of determining whether a single effect unit (e.g., a gene or SNP) is jointly associated with a specific subset of features.

A direct way to obtain subset-level claims is to use the partial conjunction hypothesis (PCH) framework [8], which tests a composite null stating that at least one feature in the subset is null; rejection then provides direct evidence that the subset is jointly significant. Ray and Chatterjee [9] clarified that this composite-null formulation is distinct from a global-null test and is better suited to pleiotropy detection.

Although many existing methods are described as subset-based approaches or cross-phenotype association tests, they can often be understood, either explicitly or implicitly, within the PCH framework [6, 9–13]. A common strategy in this framework is to assign each effect unit a binary significance pattern across features and to learn the frequencies of these patterns together with a pooled non-null effect-size distribution. Existing implementations often rely on relatively simple assumptions for the marginal effect-size distribution, and can become anti-conservative or overly conservative when estimation errors are correlated. These limitations motivate APCH-b, which retains binary configurations while using adaptive shrinkage (ash) [14] to flexibly model the feature-wise marginal effect-size distributions and to explicitly account for estimation error correlation.

For PCH, pooling all active effects of a feature into a single marginal prior can also be restrictive. Joint signals may be driven by a shared biological mechanism, such as a common pathway, so a SNP need not behave the same way when it is associated with only one trait as when it appears in a jointly associated subset. This motivates APCH-s, which enriches the binary significance pattern with latent magnitude states and thereby allows configuration-specific magnitude sharing. Finally, in multivariate settings, exact zeros can be difficult to distinguish from weak nonzero effects, and sub-null configurations, where only a subset of features is non-null, can be difficult to distinguish from joint non-null configurations. We therefore introduce the subset-level local false sign rate (lfsr) in addition to the subset-level local false discovery rate (lfdr), and find that the lfsr often provides a more robust inferential criterion.

In this article, we propose APCH (Adaptive Partial Conjunction Hypothesis). In simulations, APCH remains powerful under heterogeneous effects, with well-calibrated error control. We illustrate APCH with a meta-analysis of five type 2 diabetes (T2D)-related traits. T2D exhibits substantial genetic heterogeneity and the underlying molecular mechanisms remain incompletely understood [15, 16]. We use APCH to identify co-occurring trait patterns and candidate loci. To probe mechanisms underlying these loci, we then apply APCH to a multi-tissue transcriptome-wide association study (TWAS) by integrating GWAS summary statistics with trained gene-expression prediction models across multiple tissues [17, 18]. APCH is implemented in an R package available on [GitHub](https://github.com/colinyuxin/APCH).

## 2 Method

### 2.1 Method overview

We work with summary statistics from association studies: each SNP or gene is an effect unit and each trait or tissue is a feature. An overview of APCH is shown in Figure 1.

**Figure 1.**
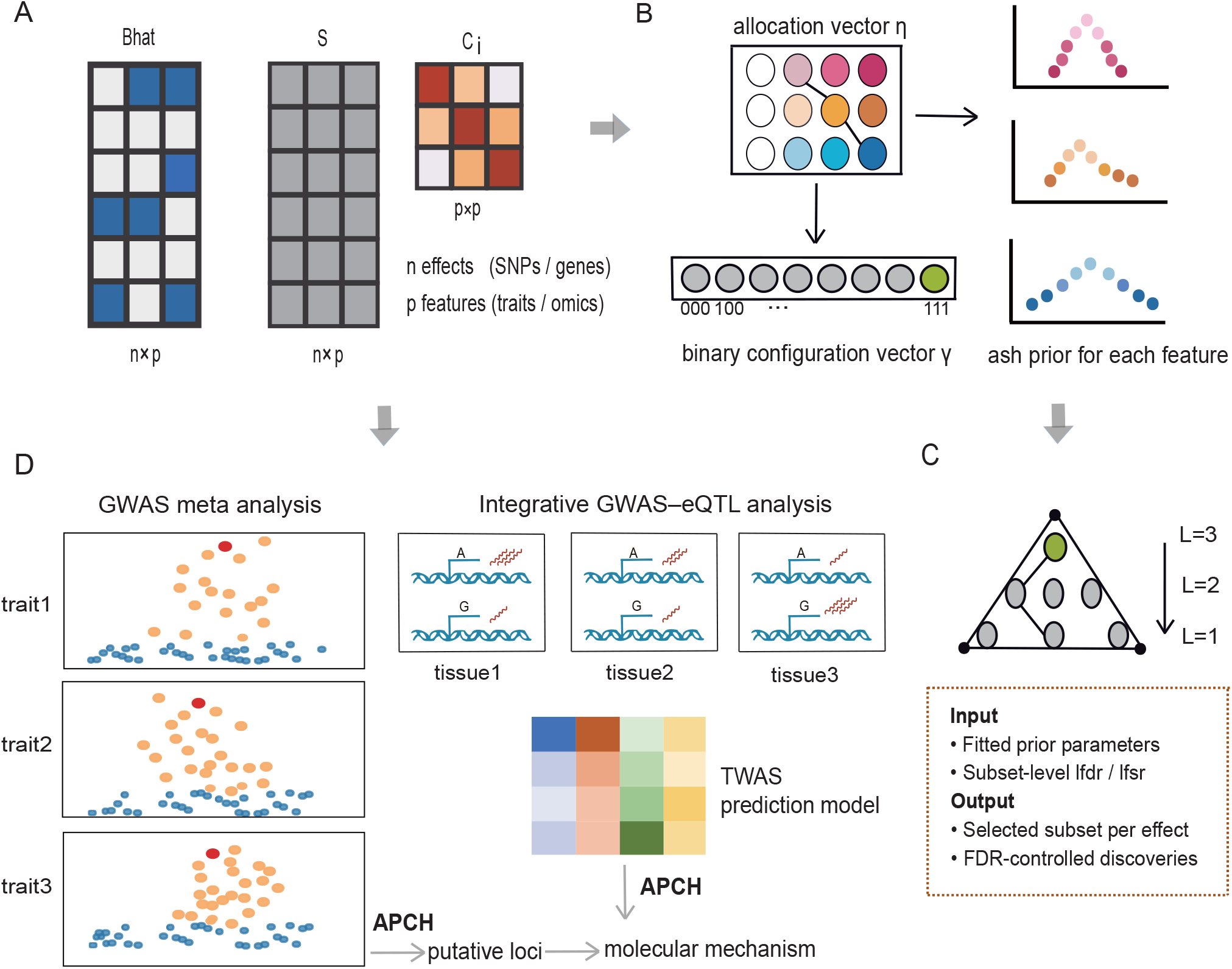
Graphic overview of APCH. (**A**) APCH takes multi-feature summary statistics as input: effect estimates 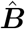, standard errors ***S***, and estimation error correlations ***C***_*i*_. (**B**) The main APCH-s model uses a hierarchical shrinkage prior: an allocation vector ***η***_*i*_ selects grid components for each feature’s ash prior and induces a binary configuration ***γ***_*i*_ that records the null/non-null pattern. (**C**) APCH makes subset-level discoveries, where a reported subset is false if any included feature is null. Candidate subsets are searched from larger to smaller sizes using subset-level lfdr or lfsr. (**D**) Example applications: cross-trait meta-analysis and integrative GWAS–eQTL analysis using TWAS prediction models for multiple tissues.

Index the *p* features by *j* = …, *p*, and let *U* = {*U ⊆* {1, …, *p*} : |*U*| ≥ 1} denote all non-empty feature subsets. Let *b*_*ij*_ (*i* = 1, …, *n*; *j* = 1, …, *p*) denote the true value of the *j*-th feature of the *i*-th effect (e.g., the effect size of the *i*-th genetic variant on the *j*-th trait).

For each effect *i* and subset *U*, we consider the partial conjunction hypotheses:

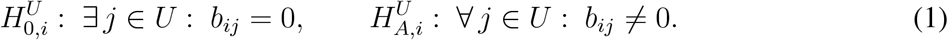

A global-null test can be driven by a single feature, so pleiotropy calls for PCH that separates partial-significance patterns from simultaneous multi-feature signals [9]. We therefore call a reported set *Û*_*i*_ a false discovery if it contains any truly null feature.

We develop a flexible hierarchical empirical Bayes (EB) model for the PCH problem. EB methods are particularly attractive in modern GWAS, where large numbers of effects and increasing sample sizes allow prior structure to be learned efficiently from the data [19]. Our guiding idea is (1) to learn the information that is essential for PCH inference and (2) to introduce sharing structure that aligns with the PCH inference goal. This leads to a nested family of two models. We start from APCH-b, which learns from the data the frequencies of all significance patterns and the feature-wise marginal effect-size distributions. This idea extends the classical two-group model by lifting the single-feature null/non-null indicator to a binary configuration across *p* features. Extensions of the two-group model for PCH have been explored before [6, 10, 11], while our key distinction is to model the feature-wise marginals with ash [14], a flexible data-adaptive shrinkage prior that captures complex genetic architecture. APCH-s extends APCH-b by modeling cross-feature magnitude sharing, which is essential when the effect-size pattern among active features depends on the underlying configuration. APCH-s involves a substantially larger parameter space and is therefore more computationally demanding. To improve computational efficiency, we adopt an almost-free pruning strategy that exploits the nested structure of PCH and substantially reduces the parameter space. Finally, we use a simple level-by-level rule for selecting subsets based on subset-level posterior quantities. In particular, we define subset-level lfdr and lfsr and recommend lfsr as the primary inferential quantity.

### 2.2 Mathematical description of APCH

#### 2.2.1 Model setup and assumptions

Let *b*_*ij*_ (*i* = 1, …, *n*; *j* = 1, …, *p*) denote the true value of the *j*-th feature of the *i*-th effect, 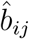 the corresponding observed estimate, and *s*_*ij*_ the standard error. Let ***b***_*i*_ = (*b*_*i*1_, …, *b*_*ip*_) and 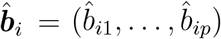. APCH is formulated as an empirical Bayes multivariate normal means (EBMNM) model [20, 21]. The EBMNM model assumes that we observe vectors 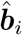 that are independent, noisy, normally distributed measurements of underlying true values ***b***_*i*_:

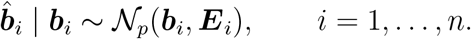

The covariances ***E***_*i*_ are specified as: ***E***_*i*_ = ***S***_*i*_ ***C***_*i*_ ***S***_*i*_. Here ***S***_*i*_ is diagonal with entries *s*_*i*1_, …, *s*_*ip*_, and ***C***_*i*_ is the *p ×p* correlation matrix of estimation errors across features for effect *i*. ***E***_*i*_ are assumed to be known and positive definite. The EBMNM model further assumes that the unknown means are independent and identically distributed draws from some distribution 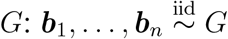. The goal is to learn posterior quantities induced by *G*, such as the lfdr. For convenience, we work on the *z*-score scale throughout, so that 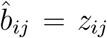 and *s*_*ij*_ = 1. Our model captures within-effect noise correlation across features, which in GWAS may arise from sample overlap [22] or kinship [23].

#### 2.2.2 APCH-b as a natural extension of the two-group model

In the two-group model, each effect unit is either null or non-null, and we estimate the non-null effect-size distribution *g*(*·*) together with the non-null proportion from the data. For *p* features, we generalize this to a binary configuration vector ***γ***_*i*_ ∈ {0, 1}^*p*^ and assign a prior Pr(***γ***_*i*_ = ***r***) = *ϕ*_***r***_ for ***r*** ∈ {0, 1}^*p*^. This yields 2^*p*^ significance patterns, corresponding to all possible partial conjunction hypotheses across the *p* features. The resulting hierarchical model can be written as:

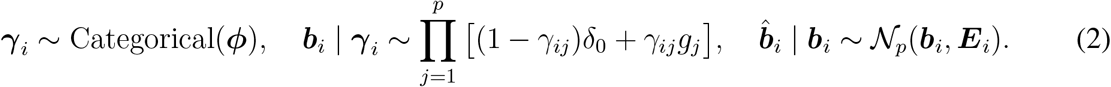

Here *δ*_0_ denotes a point mass at zero. In practice, we use a two-stage strategy: first estimate the feature-wise non-null marginals *g*_*j*_, then estimate the configuration frequencies *ϕ*_***r***_ within this model. Our key choice is to estimate *g*_*j*_ via adaptive shrinkage (ash), which approximates an arbitrary uni-modal distribution by a scale mixture of centered normals:

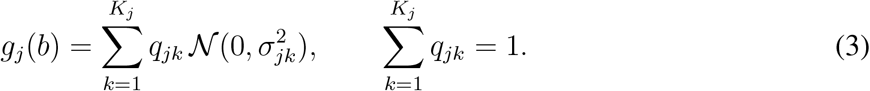

Here {*σ*_*jk*_} is a feature-specific grid of scales and *q*_*jk*_ are the mixture weights.

#### 2.2.3 APCH-s enabling configuration-specific magnitude structure

APCH-b learns the frequencies of binary significance patterns while pooling the magnitude of effects into a feature-wise marginal prior. APCH-s lifts the prior from binary configurations ***γ*** to latent magnitude states ***η***, thereby allowing configuration-specific magnitude structure.

Specifically, APCH-s introduces an allocation vector ***η***_*i*_ = (*η*_*i*1_, …, *η*_*ip*_), where *η*_*ij*_ = 0 indicates that feature *j* is null for effect *i*, and *η*_*ij*_ = *k* ∈ {1, …, *K*_*j*_} indicates that feature *j* is assigned to the *k*-th non-null magnitude state, that is, the *k*-th ash grid for feature *j*. The corresponding binary configuration is induced by *γ*_*ij*_ = **1**{*η*_*ij*_ > 0}. The resulting hierarchical model is

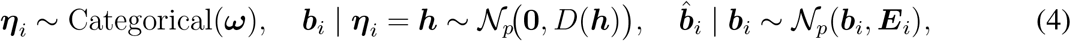

where 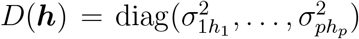, with the convention that 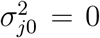 under the exact-null model. The feature-wise grids {*σ*_*jk*_}_*k*≥1_ are still treated as fixed plug-in inputs estimated from the marginals, serving as a feature-specific dictionary of representative effect magnitudes. Under APCH-s the allocation weights *ω*_***h***_ are learned freely. APCH-b is recovered as the restricted special case in which 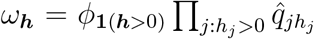, where 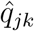 denotes the estimated feature-wise conditional mixture weight from the marginal fit.

#### 2.2.4 Posterior quantities and inference

We define the subset-level local false discovery rate as

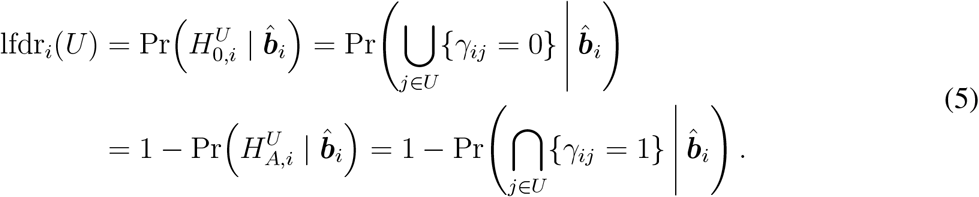

In words, lfdr_*i*_(*U*) is the posterior probability that any feature in *U* is null, or equivalently one minus the posterior probability that all features in *U* are simultaneously non-null. This also makes clear that sub-null configurations complicate PCH because they must be distinguished from joint non-null effects. We further define the subset-level local false sign rate as

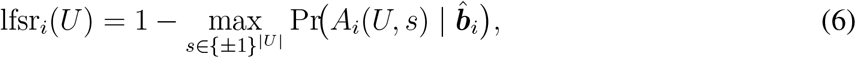

where *A*_*i*_(*U, s*) := ⋂_*j*∈*U*_ {*s*_*j*_*b*_*ij*_ > 0} denotes the event that all effects in subset *U* follow the joint sign pattern *s*. Thus, compared with the lfdr, the lfsr imposes double shrinkage: it favors effects that are jointly non-null on *U* and further shrinks those whose joint sign pattern is unstable. One may therefore expect truly jointly significant effects to be only mildly shrunk.

Let *ℋ*_*U*_ = {***h*** : *h*_*j*_ > 0 for all *j* ∈ *U*} denote the set of allocation states that are active on all features in *U*, and *ℬ*(*U, s*) = {***b*** ∈ ℝ^*p*^ : *s*_*j*_*b*_*j*_ > 0 for all *j* ∈ *U*} the region in which the effects in *U* follow the joint sign pattern *s*. Then for our model:

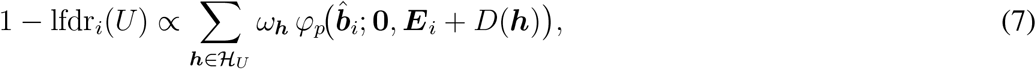

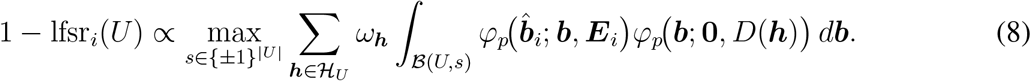

Here *φ*_*p*_(*·*; *µ*, Σ) denotes the *p*-variate Gaussian density. Because the exact null can be difficult to distinguish from weak nonzero signals in multivariate settings, we recommend the lfsr as the primary inferential criterion, as it is often more stable and scientifically interpretable [14].

We obtain these posterior quantities by Bayes’ formula, estimate the unknown model parameters by maximum likelihood, and plug the resulting estimates into the posterior formulas to obtain 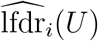 and 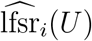. Moreover, if *U*_1_ *⊆ U*_2_, then lfdr_*i*_(*U*_1_) *≤* lfdr_*i*_(*U*_2_). APCH thus performs subset selection through the following level-by-level procedure, moving from larger to smaller subset sizes:

1. At level *L*, for each effect not yet selected, retain its best size-*L* candidate, the subset with the smallest estimated lfdr/lfsr among all subsets of size *L*.
2. Rank these candidates by increasing lfdr/lfsr, and accept them sequentially as long as the running average of the accepted lfdr/lfsr does not exceed *α*.
3. Once an effect is accepted, remove it from subsequent lower-level searches.

Level-by-level inference can be viewed as a multivariate extension of *q*-value-based FDR control [24].

#### 2.2.5 Parameter estimation and allocation-state pruning

Once the feature-specific hyperparameters are fixed, both APCH-b and APCH-s can be estimated within a common computational framework as finite mixture models with known component densities and unknown mixture weights. We estimate these weights by maximizing a weakly regularized likelihood using mixsqp [25]. We place a weak Dirichlet prior on the allocation weights, which can reduce estimation variance while introducing little bias. Moreover, all required convolutions between the prior components and the Gaussian observation model admit closed-form expressions by Gaussian conjugacy, which makes the computation efficient.

For APCH-s, the full allocation space can be large, so we further stabilize and accelerate estimation by pruning unsupported allocation states before the final joint fit. The key observation is that PCH is nested across subsets. For example, if a full binary configuration is 11101 and we consider the subset *U* = {1, 3, 5}, then the corresponding local configuration is 111. The weight of a local configuration can be obtained by summing the weights of all full configurations that agree with it on that subset, that is, by marginalizing out the remaining coordinates. If a local configuration has zero weight, then every compatible full configuration must also have zero weight. The same logic also applies to allocation states. Exact testing of zero-weight, potentially overfitted, components in a finite mixture model can be carried out by a parametric refit bootstrap [26, 27], although this is computationally infeasible here. Our goal is to prune allocation states that have negligible impact on inference. We fit pairwise and three-way APCH-s models and remove local allocation states with little posterior support. After a local state is pruned in a low-order subproblem, every full allocation state compatible with that local state is removed from the final APCH-s support. For example, if a pairwise state is pruned on features (*j, k*), then any full allocation state with the same entries on (*j, k*) is removed. Details are provided in the Supplementary Notes.

### 2.3 Simulation design

#### Simulation Study 1

Simulation Study 1 provides a baseline for evaluating APCH-b and existing methods for subset-based inference under the PCH framework. Effects are conditionally independent across traits given the binary configuration. Within this setting, we vary the marginal effect-size distribution and the correlation of estimation errors to assess robustness to non-Gaussian marginals and correlated noise. We compared APCH with OPERA [6], RepFDR [10], PLACO/PLACO+ [9, 13], and CPBayes [11].

#### Simulation Study 2

The competing methods in Simulation Study 1, either explicitly or implicitly, adopt an APCH-b–type paradigm by pooling active effects within each feature and overlooking heterogeneity across effect units. This simplification reduces model complexity, but may miss important sharing structure in realistic data.

We therefore compare APCH-b and APCH-s under different sharing structures in Simulation Study 2. We also compare with Mash [20, 21], a well-established EBMNM model that learns sharing patterns from the data. Although Mash was not designed for the PCH problem, we compute the same subset-level lfsr for a fair comparison. We generate the data from structural equation models, motivated by the idea that pleiotropy may be driven by a combination of shared latent factors and trait-specific signals. This view has been widely adopted in genetic studies [28, 29].

#### Simulation Study 3

Simulation Study 3 examines how the strength of sharing within the joint non-null configuration affects APCH-s. We fixed the proportions of global null, partial-significance, and joint non-null effects. The four scenarios differed only in the sharing structure of effects within the joint non-null configuration: conditional independence given the binary configuration, corresponding to the APCH-b regime; conditional independence given a shared allocation vector, corresponding to the APCH-s regime; and moderately or totally correlated joint non-null effects, which are misspecified relative to the APCH-s model.

#### Evaluation metrics

For a reported subset *Û*_*i*_, let 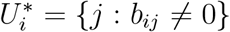 denote the true active subset. We report the FDR, under which a reported subset is counted as a false discovery if 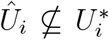. We also report the false sign rate (FSR). Under this criterion, a reported subset is counted as correct only if 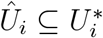 and 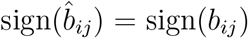 for all *j* ∈ *Û*_*i*_. Thus, among reported subsets with 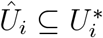, we distinguish true-sign discoveries, for which all reported signs are correct, from boundary discoveries, for which at least one reported sign is incorrect. The FSR asks whether the posterior decision agrees with the true effect direction. Intuitively, it is unsatisfactory to claim a significant discovery when the reported signal direction has, in fact, been reversed by noise. Under this classification, FDR counts only false discoveries as errors, whereas FSR counts both boundary and false discoveries as errors. Detailed data-generating procedures and implementation details for competing methods are provided in Supplementary Note S2.

### 2.4 Real data analysis

#### 2.4.1 Meta-analysis of five T2D-related traits

Unless otherwise stated, APCH refers to the pruned APCH-s model. We analyzed GWAS summary statistics for five traits related to T2D: type 2 diabetes (T2D), fasting insulin adjusted for BMI (FI), fasting proinsulin adjusted for BMI (PROI), fasting plasma glucose (FPG), and 2-hour post-challenge glucose (2hPG) [30–32]. FI and PROI serve as proxies for two major etiologic pathways of T2D, insulin resistance and beta-cell dysfunction, respectively. FPG and 2hPG quantify complementary dimensions of glucose metabolism, capturing fasting hepatic glucose regulation and post-challenge glucose handling.

We kept SNPs that were observed in all five traits. Selecting the SNP universe for APCH is a challenging step, because the input variants determine which loci enter the empirical Bayes model and can therefore affect the inferred cross-trait patterns. Our goal was to select representative variants for each locus while minimizing selection bias. We considered two main sources of bias. First, LD can distort the representation of loci: if many correlated SNPs are retained, loci with denser or more complex LD structure may be overrepresented; if pruning is too aggressive, some loci may lose useful representatives. Second, SNP selection may also be biased if it is driven either by one particular trait or by a specific cross-trait summary, such as an omnibus test that aggregates evidence across traits; in either case, the resulting input set may be enriched for the corresponding trait or association pattern.

For the primary analysis, we used a random LD-thinning strategy based on predefined UKBB-LD regions and an LD threshold of *r*^2^ = 0.25. Within each region, variants were selected according to their first occurrence in the predefined UKBB-LD ordering. This random-thinning idea is similar to the strategy used by Urbut et al. [33]. We further compared alternative SNP-selection strategies as sensitivity analyses. These included LD thinning under different thresholds (*r*^2^ = 0.1, 0.25, 0.6), single-trait lead-SNP selection using each trait as the anchor, and omnibus-test-based lead-SNP selection using MetaUSAT [34] *p*-values. The corresponding results are reported in the Supplementary Notes (Supplementary Tables 2–5).

To summarize the findings at the locus level, we defined tag SNPs within the LD blocks induced by the *r*^2^ *<* 0.25 clumping, choosing as the tag the SNP with (i) the largest number of traits for which APCH declared joint significance and (ii) the smallest corresponding lfdr/lfsr within that block. We then constructed finer-scale LD loci by reclumping these tag SNPs using *r*^2^ *<* 0.01 as the within-locus threshold. SNP-to-gene mapping for reported loci was performed using FUMA [35].

#### 2.4.2 Joint analysis of multi-tissue TWAS

To link T2D loci to gene expression, we applied APCH to a multi-tissue TWAS of T2D. We used GTEx v8 elastic-net expression prediction models [36] for six relevant tissues: subcutaneous adipose (SAT), visceral adipose (VAT), liver, skeletal muscle, pancreas, and whole blood. Single-tissue *z*-scores were computed with S-PrediXcan [18]. Because the predicted expression in each tissue is a linear function of overlapping sets of SNPs, the estimation errors are correlated across tissues, and we took this correlation from S-MultiXcan [37]. Following Liang et al. [38], we calibrated the *z*-scores for the null inflation caused by polygenicity.

Data sources and processing details are provided in Supplementary Note S4.

## 3 Results

### 3.1 Simulation Studies

#### Comparison with existing PCH methods

Inspired by the direct-pleiotropy setting in GWAS-Brewer [39], we generated two-trait summary data for 10,000 independent effects, with effect-size distributions ranging from normal to spiky, fixed noise scale, and weak-to-strong noise correlation *ρ*. We considered both a mixed setting with sub-null and joint non-null signals and a restricted setting containing only global-null and joint non-null configurations with varying global-null proportion. Figure 2 summarizes the results.

**Figure 2.**
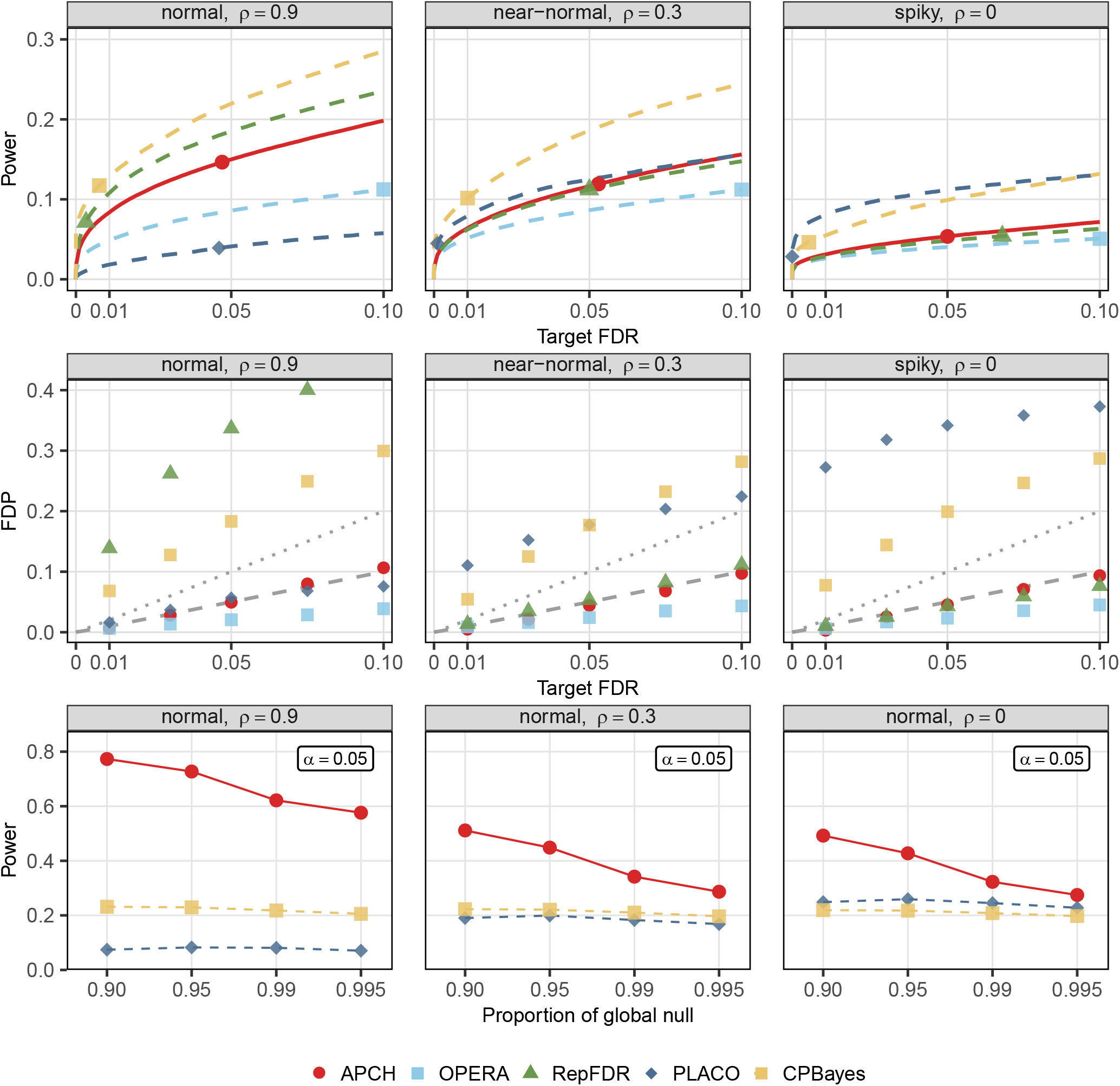
Calibration and power under baseline PCH settings. Simulation Study 1 varies three factors: the marginal effect-size distribution, the estimation-error correlation *ρ*, and the abundance of joint signals. The top two rows correspond to the mixed configuration setting (*φ*_00_, *φ*_10_, *φ*_01_, *φ*_11_) = (0.80, 0.10, 0.05, 0.05). The bottom row corresponds to the setting with only global-null and joint non-null configurations, with the proportion of global null varying from 0.90 to 0.995. Power and FDP are calculated for joint non-null effects. The top row marks the points at *α* = 0.05. The second row includes reference lines *y* = *x* and *y* = 2*x*. Results are averaged over 100 replicates. FDP, false discovery proportion.

Simple Gaussian models are often insufficient to describe GWAS effect-size distributions [40, 41]. The effect-size distributions in the top and middle rows have nearly the same second moment, 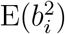, but differ in higher moments, ranging from normal to spiky. The competing methods were more sensitive to these departures from Gaussian marginals, whereas APCH remained well calibrated across settings. Performance was also affected by noise correlation: APCH gained power as *ρ* increased while maintaining calibration.

The power of EB methods depends on the abundance of signal in the data. The bottom row shows that even when the joint non-null proportion was only 0.5%, APCH still retained substantial power. Comparing the mixed and restricted settings further shows that the presence of sub-null configurations [9] makes exact joint non-null detection harder and reduces power overall. In these simulations, APCH-b and APCH-s performed similarly (Supplementary Figure 1), and both achieved higher power than the competing methods while remaining well calibrated.

#### Comparison with Mash under structural equation models

Figure 3 and Supplementary Figure 2 summarize Simulation Study 2. APCH-b performs well in Scenario 1, but deteriorates once abundant marginal signals are introduced. In our design, these marginal effects are individually weak relative to the structured joint factors, yet they are numerous. This setting is particularly relevant to GWAS, where a large fraction of effects are small and joint significance may plausibly be driven by a few shared factors. Because APCH-b pools non-null effect sizes at the feature level, it can become overly optimistic about joint significance patterns when weak marginal signals dominate the feature-wise non-null distribution.

**Figure 3.**
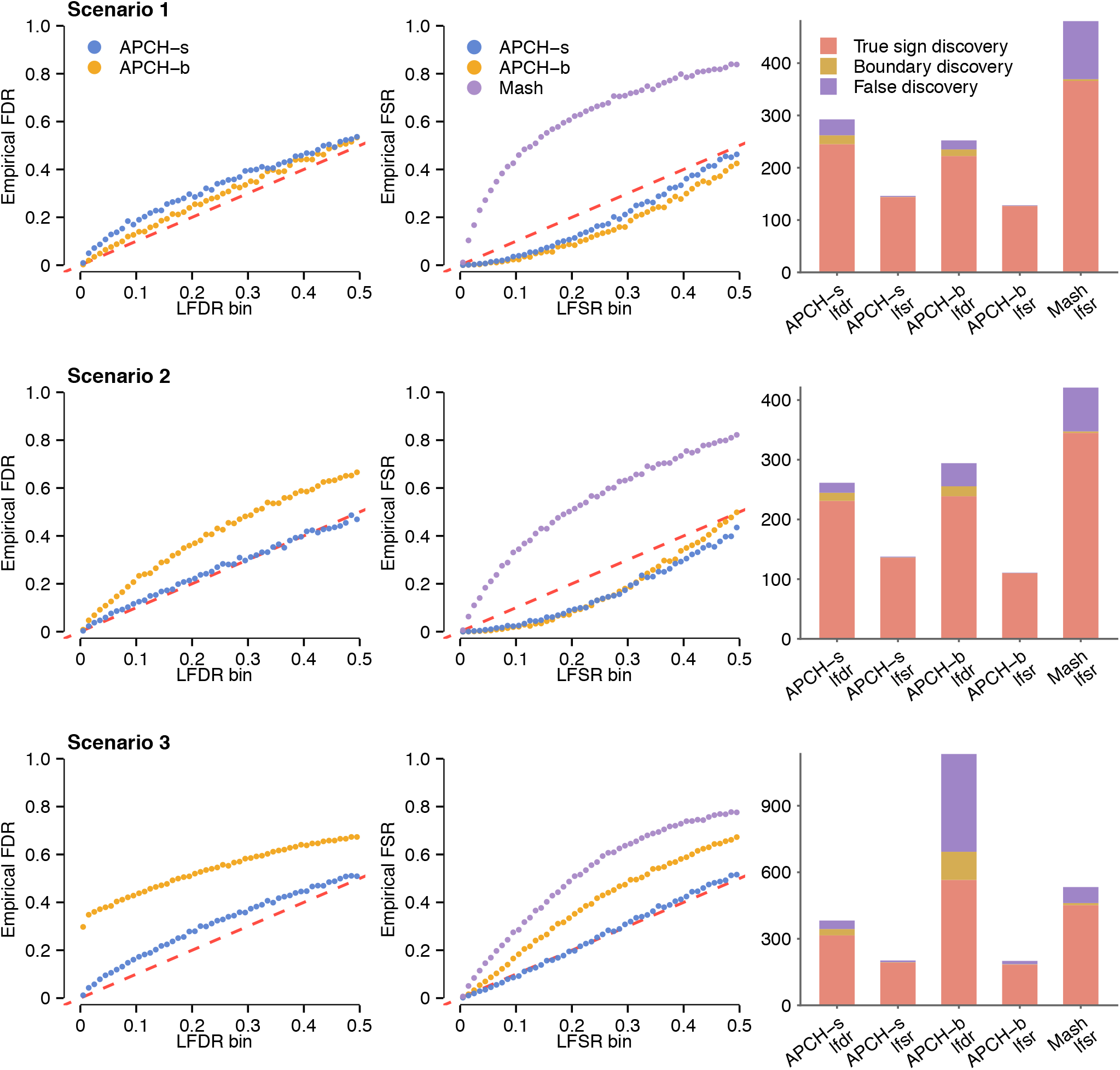
Comparing APCH-s, APCH-b, and Mash under structural equation models. Simulation Study 2 compares APCH-s, APCH-b, and Mash under three scenarios. The scenarios form an increasing ladder of difficulty, moving from a clean two-factor structure, to the presence of abundant marginal signals, and then to overlapping latent factors. The first two columns compare empirical FDR and FSR with binned lfdr and lfsr, respectively, using 50 bins over the interval [0, 0.5]; well-calibrated posterior quantities should lie close to the diagonal reference line. The third column shows the average number and composition of reported joint non-null discoveries, classified as true-sign discoveries, boundary discoveries, or false discoveries. Results are averaged over 500 replicates.

APCH-s performs well overall across all three scenarios and is only slightly anti-conservative under lfdr-based inference in Scenarios 2 and 3. This reflects an intrinsic difficulty of multivariate PCH inference: once noise acts across several coordinates, it becomes easier for weak or nearby configurations to resemble a jointly non-null pattern. This appears in two ways. First, some effects have the correct true configuration but at least one sign is flipped by noise, yielding boundary discoveries. Second, effects from nearby configurations can also be pushed toward the target configuration. This is one reason we introduce the subset-level lfsr, which effectively removes most boundary discoveries and also reduces errors arising when nearby configurations are pushed toward the target configuration by noise. Another reason will be illustrated in the real-data analysis.

Mash models effect sizes through a continuous multivariate prior, which can make it harder to sharply separate noise from weak signals and therefore harder to identify distinct configurations. We also observe much larger variance in the boxplots (Supplementary Figure 2) for Mash, partly because its performance depends on whether the fitted model successfully captures the true sharing pattern in each replicate. In our simulations, this leads Mash to overestimate signals, most noticeably in Scenario 1, where the true configurations are concentrated at subset levels 3 and 5.

These results highlight two points. First, APCH-b can be viewed as a pooling approximation to APCH-s, in which active effects are pooled into feature-wise marginal priors. This approximation can become unstable when the data contain abundant but weak marginal signals or configuration-specific effect-size patterns, supporting APCH-s as our main model. Second, lfsr-based inference improves the quality and stability of subset discoveries. In Scenarios 2 and 3, APCH-b is highly anti-conservative under lfdr-based inference. Switching to lfsr restores error control, although the resulting calls become conservative. This is because lfsr adds a sign-aware layer of shrinkage and filters out many boundary discoveries. However, lfsr alone is not intrinsically tied to the PCH testing problem and does not guarantee error-rate control.

#### Robustness to model misspecification

Genetic correlation [42] provides another perspective for understanding pleiotropy. It focuses on pairs of traits that have similar directions of effects across different SNPs. For APCH, however, we do not model the correlation of effects. Across the four scenarios, the two traits have genetic correlations *r_g_* = 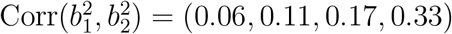, but squared-effect correlations of 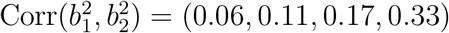. Only the latter matters for APCH, whose priors are symmetric around zero. The second moment of each trait, 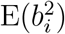, is the same across the four scenarios. Figure 4 C illustrates the observed effect distributions under different sharing structures. As the degree of sharing increases, APCH power also improves. Although APCH does not model the effect correlation in Scenarios 3 and 4, it still benefits from it, because correlated effects also share magnitudes, which shared allocation states capture. Consistently, the proportion of true discoveries with both trait *p*-values below 10^*−*4^ rose from 0.62 to 0.64 and 0.76 from Scenario 2 to Scenario 4 (Figure 4 A). APCH is only mildly anti-conservative at large *α* in the fully correlated scenario (Figure 4 B). These results suggest that modeling shared allocation states captures most of the useful structure for PCH inference; explicit modeling of effect correlation may matter mainly when the correlation is very strong. APCH-b, of course, cannot capture different sharing structures, and thus is more sensitive to model misspecification. It is also notable that the marginal distribution is the same across different configurations. Thus, the misspecification here comes mainly from the sharing structures, not from the pervasive but moderate to weak effects as in Simulation Study 2. We compared APCH-b and APCH-s in Supplementary Figure 3.

**Figure 4.**
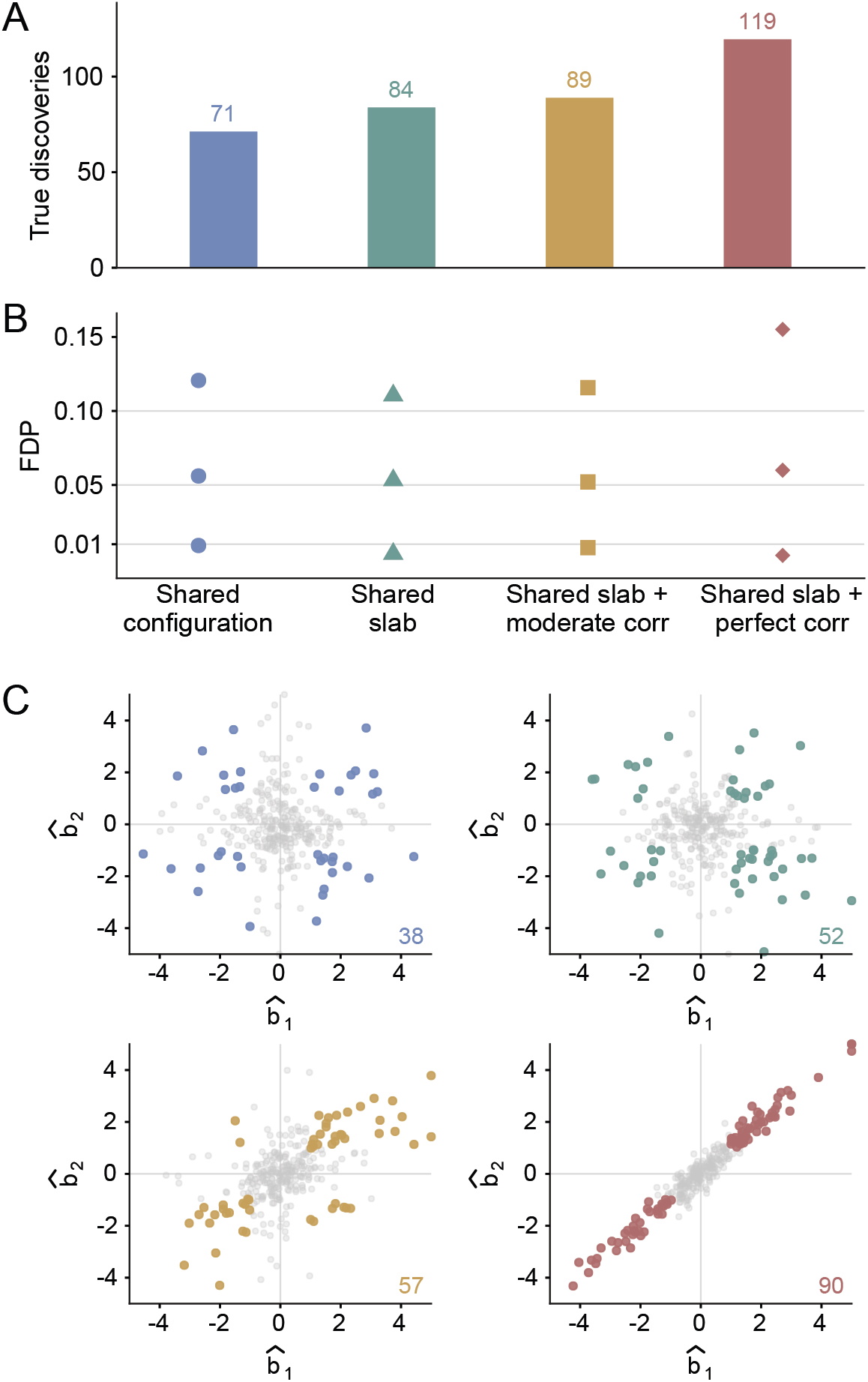
Joint discoveries by APCH-s under different effect-sharing scenarios. (A) Mean number of true joint non-null discoveries using APCH-s with lfsr-based inference at *α* = 0.05 across 100 replicates. (B) FDP of joint non-null effects across target FDR levels for the four effect-sharing scenarios, using APCH-s with lfsr-based inference. (C) Example observed effect estimates 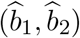 for joint non-null effects under each scenario. Colored points denote effects for which both marginal *p*-values are below 10^*−*4^; numbers give the corresponding counts.

### 3.2 Real data analysis

#### Joint signals are not rare, but lfsr removes near-zero partners

Before presenting the APCH discoveries, we first examine the difference between subset-level lfdr and lfsr. In both the GWAS and the TWAS, lfsr removed part of the discoveries declared jointly significant by lfdr, and more so in the GWAS (Figure 5B, D). Two features explain this. First, in PCH inference the joint non-null configuration is not necessarily rarer than a partial-significance configuration. With an anchor trait such as T2D, a variant affecting fasting glucose but not T2D is more surprising than one associated with both, and in the TWAS, eQTLs are widely shared across tissues [20, 36]. Once one trait has strong evidence, the posterior odds of the joint configuration versus the configuration in which only that trait is non-null equal their prior odds times the Bayes factor. In our fits these prior odds favor the joint configuration, so weak evidence for the second trait is enough for a small lfdr. This is the opposite of single-trait inference, where the prior favors the null. A related effect is known in classical EB multiple testing, where a small estimated null proportion can make the FDR threshold less stringent than the nominal single-test threshold [43, 44]. Second, the GWAS contains many moderate-to-null signals, so the marginal ash fits place most non-null mass on small slabs, except for PROI, whereas the TWAS fits have no such slabs (Supplementary Table 7). An effect near the boundary can then be explained by either a small slab or an exact zero, which lfdr cannot separate, whereas lfsr also requires a stable sign. Therefore, lfdr-based inference is not necessarily incorrect, but lfsr-based inference gives a more robust and more interpretable set of discoveries.

**Figure 5.**
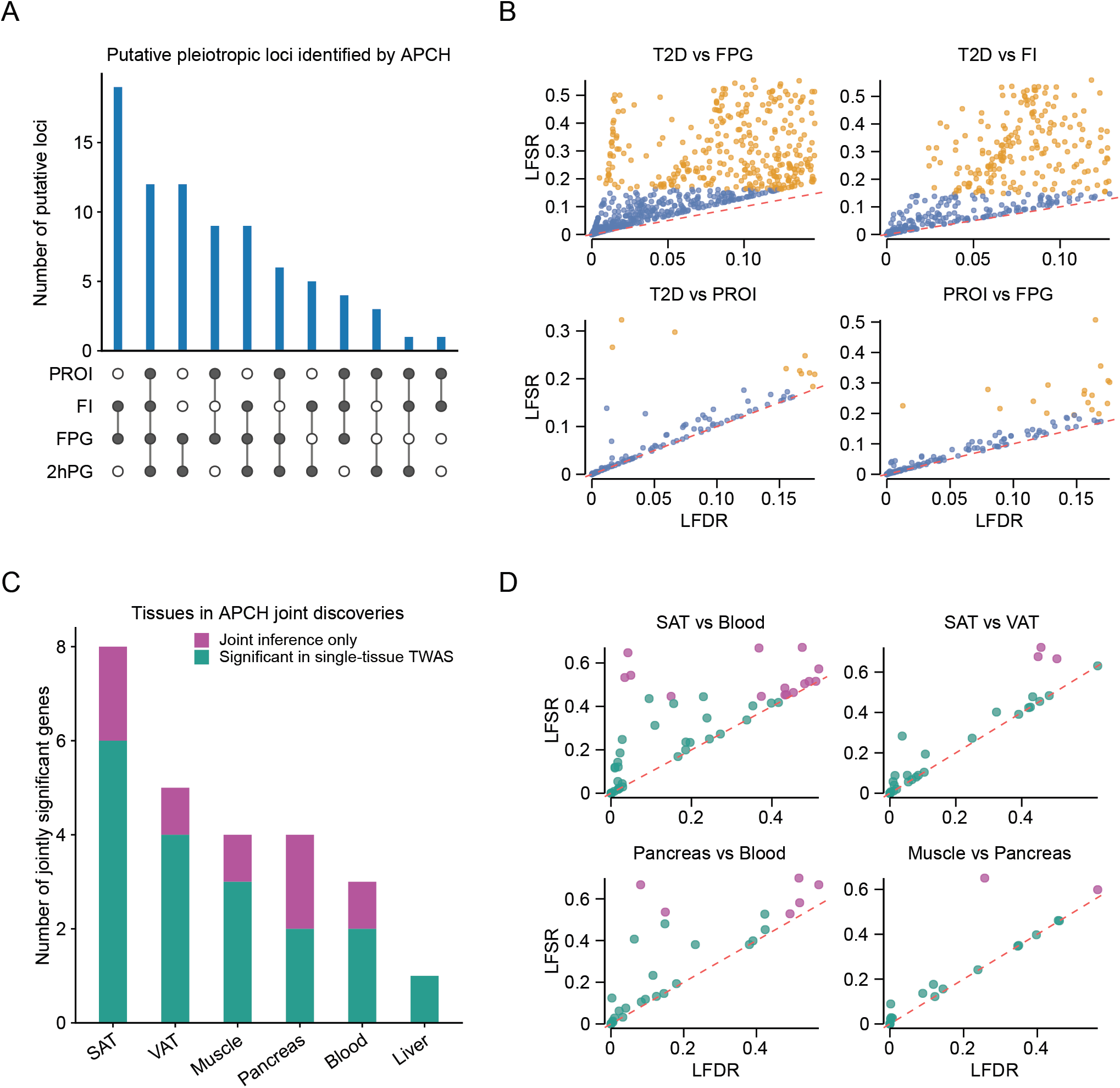
Joint discoveries in T2D-related GWAS traits and multi-tissue TWAS. (A) Putative pleiotropic loci identified by APCH (lfsr at *α* = 0.01) for trait combinations including T2D and at least three traits. (C) Number of jointly significant genes involving each tissue (lfsr at *α* = 0.01). Green bars mark genes that are also significant in a single-tissue analysis of that tissue (ash, lfsr at *α* = 0.01). (B, D) APCH was run separately for each trait pair (B) and tissue pair (D). Each point represents an effect unit declared jointly significant for the pair using lfdr at *α* = 0.05 (B) or *α* = 0.2 (D). Points significant under both lfdr- and lfsr-based inference are blue (B) or green (D); points significant under lfdr only are orange (B) or purple (D). The red dashed line denotes *y* = *x*.

#### APCH discovers co-occurring patterns in T2D

APCH identified 81 putative pleiotropic loci associated with at least three traits by lfsr using *α* = 0.01, including 12 loci jointly significant across all five traits (Figure 5 A, Table 1). In the TWAS, APCH identified 10 genes jointly significant in at least two tissues, including *JAZF1* in five tissues (Figure 5 C, Supplementary Table 8). While colocalization [45] and fine-mapping [46] analyses are typically used to resolve individual loci, APCH provides a genome-wide overview of how signals are shared across traits and tissues.

**Table 1.**
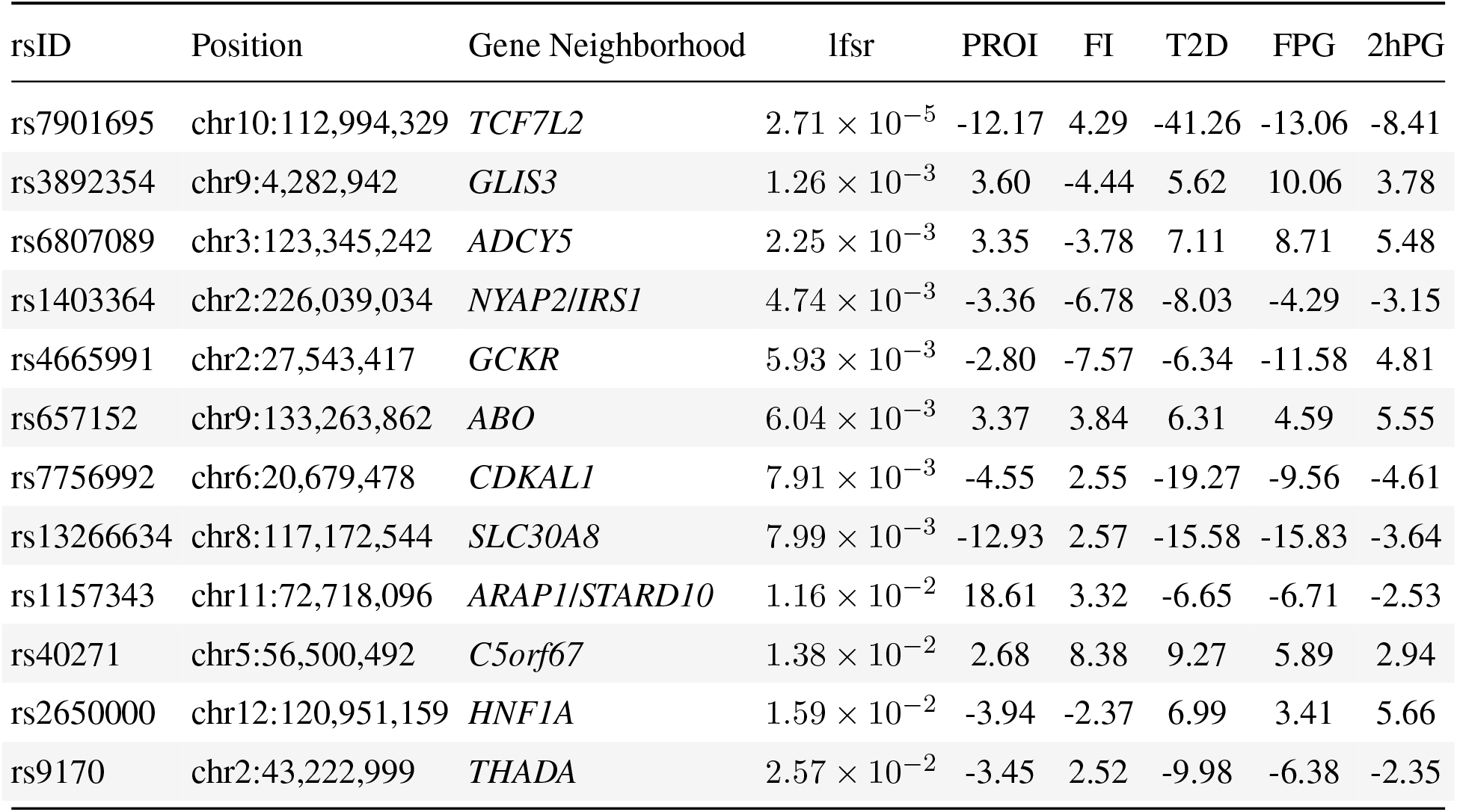
Tag SNPs identified by APCH as jointly significant for all five traits under lfsr-based inference at *α* = 0.01. Genomic positions are reported on the hg38 build. Observed *z*-scores are shown for the five traits. Combined Gene Neighborhood labels such as *NYAP2*/*IRS1* indicate multiple nearby plausible genes.

| rsID | Position | Gene Neighborhood | lfsr | PROI | FI | T2D | FPG | 2hPG |
| --- | --- | --- | --- | --- | --- | --- | --- | --- |
| rs7901695 | chr10:112,994,329 | *TCF7L2* | $2.71 \times 10^{-5}$ | -12.17 | 4.29 | -41.26 | -13.06 | -8.41 |
| rs3892354 | chr9:4,282,942 | *GLIS3* | $1.26 \times 10^{-3}$ | 3.60 | -4.44 | 5.62 | 10.06 | 3.78 |
| rs6807089 | chr3:123,345,242 | *ADCY5* | $2.25 \times 10^{-3}$ | 3.35 | -3.78 | 7.11 | 8.71 | 5.48 |
| rs1403364 | chr2:226,039,034 | *NYAP2/IRS1* | $4.74 \times 10^{-3}$ | -3.36 | -6.78 | -8.03 | -4.29 | -3.15 |
| rs4665991 | chr2:27,543,417 | *GCKR* | $5.93 \times 10^{-3}$ | -2.80 | -7.57 | -6.34 | -11.58 | 4.81 |
| rs657152 | chr9:133,263,862 | *ABO* | $6.04 \times 10^{-3}$ | 3.37 | 3.84 | 6.31 | 4.59 | 5.55 |
| rs7756992 | chr6:20,679,478 | *CDKAL1* | $7.91 \times 10^{-3}$ | -4.55 | 2.55 | -19.27 | -9.56 | -4.61 |
| rs13266634 | chr8:117,172,544 | *SLC30A8* | $7.99 \times 10^{-3}$ | -12.93 | 2.57 | -15.58 | -15.83 | -3.64 |
| rs1157343 | chr11:72,718,096 | *ARAP1/STARD10* | $1.16 \times 10^{-2}$ | 18.61 | 3.32 | -6.65 | -6.71 | -2.53 |
| rs40271 | chr5:56,500,492 | *C5orf67* | $1.38 \times 10^{-2}$ | 2.68 | 8.38 | 9.27 | 5.89 | 2.94 |
| rs2650000 | chr12:120,951,159 | *HNF1A* | $1.59 \times 10^{-2}$ | -3.94 | -2.37 | 6.99 | 3.41 | 5.66 |
| rs9170 | chr2:43,222,999 | *THADA* | $2.57 \times 10^{-2}$ | -3.45 | 2.52 | -9.98 | -6.38 | -2.35 |

For GWAS, APCH frequently selected PROI and FI together, although they proxy distinct T2D etiologic axes related to beta-cell proinsulin processing and insulin resistance. This pattern suggests a more refined insulin-related pathway at T2D loci than would be apparent from single-trait analyses alone. The inferred directions varied across loci. Discordant PROI–FI effects are consistent with impaired proinsulin-to-insulin conversion [47], while concordant effects are consistent with insulin-resistance-related beta-cell workload [48]. A second pattern involved FPG and 2hPG. Across APCH-selected subsets, FPG showed a broader and more independent pattern, with 2hPG appearing mainly alongside FPG in larger joint patterns. This asymmetry is consistent with physiological studies separating fasting hyperglycemia from post-load glucose intolerance: FPG is more directly linked to fasting hepatic glucose regulation, while post-load glucose abnormalities more often reflect combined defects in insulin sensitivity and beta-cell response [49, 50].

Regional plots showed that APCH-selected loci can have different local architectures (Figure 6): panel A spanned a broader multi-gene region with possible multiple variants or causal-gene ambiguity, while panel B showed a more localized signal in a cleaner locus, consistent with previous studies [51– 53]. For complex regions, follow-up analyses remain necessary. Additional examples include an *L* = 4 partially significant association and a non-canonical jointly significant locus near *C5orf67* (Supplementary Figure 7).

**Figure 6.**
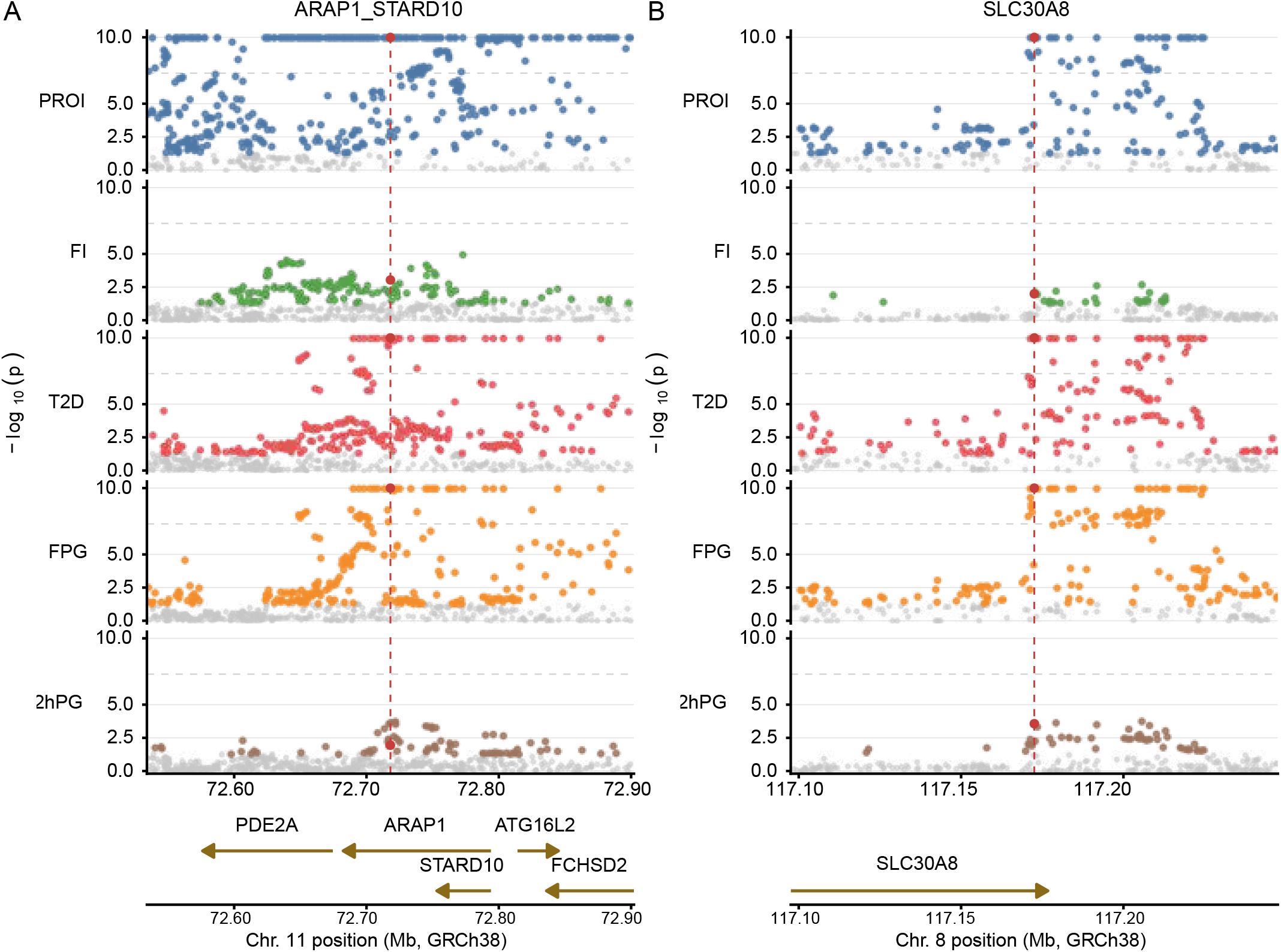
Regional association plots of joint-significant pleiotropic loci near *ARAP1*/*STARD10* (A) and *SLC30A8* (B). Shown are association *p*-values from five traits, with *−*log_10_(*p*) truncated at 10. Red points indicate the SNPs selected by APCH for prior fitting.

The jointly significant subsets often included traits whose marginal associations did not reach conventional genome-wide significance in single-trait analyses. Several “secondary” traits also showed coherent local peaks around the APCH-selected SNPs (Figure 6). Under APCH’s ash-based unimodal priors, weak nonzero effects can remain non-null after shrinkage toward zero [14]. Also, APCH aggregates evidence across different effects, so a moderate effect for one trait can still have a low lfsr when it appears in a pattern that is strongly supported by the data.

For TWAS, the data were sparser than in the GWAS. The estimated null proportion was 0.94 to 0.97 in each tissue, compared with 0.61 to 0.87 for four of the five GWAS traits (Supplementary Table 7). Even so, 7 of the 25 gene–tissue pairs in the joint discoveries fell below the single-tissue significance threshold (Figure 5 C). *IRS1*, near the *NYAP2*/*IRS1* locus in Table 1, was jointly significant in both adipose tissues with negative *z*-scores, consistent with the reduced *IRS1* expression and insulin resistance reported at this locus [48]. *JAZF1* was jointly significant in five tissues, all with negative *z*-scores, suggesting an expression effect shared across tissues. This agrees with its known role in beta-cell function under metabolic stress [54].

## 4 Discussion

We introduce APCH, an empirical Bayes framework for subset-level inference. The main difference between APCH and existing subset-based or PCH methods is that APCH models a configuration-specific prior, that is, a prior over every combination of active features. This leads to a more complex model, so we make it tractable by first plugging in the marginal variance grids as representative magnitude states and then learning only the joint sharing structure among these grid states. Simulation Studies 1 and 2 clarify this point: APCH-b improves calibration through ash-based marginal modeling, but its feature-wise pooling becomes insufficient under configuration-specific effect heterogeneity, a limitation expected for existing subset-based methods.

APCH falls within the EBMNM framework, which learns a joint distribution from noisy observed data. This perspective is useful for association studies, where the data often contain moderate to abundant signals and are based on moderate to large sample sizes. However, it seems implausible to learn the full distribution from finite and noisy data. Compared with Mash [20, 21], which aims to approximate an arbitrary distribution using sufficiently many Gaussian mixture components, our model targets a more restricted part of the distribution. Our main distinctions are twofold. First, we explicitly model binary configurations, which can be viewed as imposing strong regularization toward zero for partial-significance features. Second, we do not model correlations among the underlying true effects, which helps control model complexity.

Simulation Studies 2 and 3 support this restriction: Mash had difficulty separating joint non-null signals from partial-significance ones and APCH’s shared allocation states captured most of the PCH-relevant structure without modeling effect correlations. This distinction suggests that Mash is more suitable for estimating the main structure of data when signals are abundant and for improving effect-size estimation across different conditions. By contrast, APCH gains power and remains well calibrated when the data contain a variety of sharing structures and partial-significance configurations.

Another contribution of this work is the subset-level lfsr. Compared with lfdr, lfsr additionally requires posterior stability of the joint sign pattern, and our results show that it can produce more stable and interpretable discoveries. Moreover, lfsr can be defined under any generative Bayesian model, so the subset-level lfsr introduced here may be useful beyond APCH. In genetic studies, effects are often measured across multiple related features, such as traits, tissues, or cell types, where the same interpretation of sign-aware subsets remains meaningful. A more detailed connection between classical PCH and sign-aware inference is provided in Supplementary Note S1.

In our analysis of T2D-related traits, many traits appearing in APCH-selected jointly significant subsets did not reach genome-wide significance in single-trait scans. For example, at loci such as *TCF7L2*, a strong PROI effect is accompanied by a weaker FI association. These secondary traits would typically be discarded by genome-wide thresholds, whereas APCH retains them when they are supported by the joint posterior evidence. Subset-level inference is therefore not necessarily more stringent than univariate inference. Only one variant near *TCF7L2* reached genome-wide significance in all five traits in the full GWAS summary statistics, and only one variant had *p*-value *<* 10^*−*4^ in all five traits in our primary analysis (Supplementary Table 1). The multi-tissue TWAS showed the same pattern. Because eQTLs are widely shared across tissues, strong evidence in one tissue helped recover associations in other tissues that fell below the single-tissue threshold.

APCH is not without limitations. First, it detects statistical associations of effects among features, commonly called statistical pleiotropy [1]. However, APCH does not by itself distinguish biological pleiotropy, mediated effects, design-induced artifacts, or spurious signals arising from linkage disequilibrium between distinct causal variants in nearby genes. Second, APCH is designed for a moderate number of features (e.g., *p ≤* 8), because it must model the 2^*p*^ joint on/off configurations and becomes computationally expensive as *p* grows. Scalable subset-search methods such as fastASSET [55] are better suited to tens or hundreds of traits, but they do not directly quantify the probability that a specific subset is jointly non-null; this remains the distinctive value of APCH. Third, EB methods are often criticized for overlooking uncertainty in parameter estimation compared with a fully Bayesian approach. This can be mitigated by adopting lfsr, which is less sensitive to the fitted prior [14]. However, exploring more efficient EB models that can learn the sharing structure from finite data remains an important direction.

Notwithstanding these caveats, APCH provides a more resolved view of complex traits by turning multi-feature association signals into jointly significant subsets, and is flexible enough to capture heterogeneous sharing structures, thus supporting more robust and powerful subset-level inference.

## 5 Data and Code Availability

Summary statistics for type 2 diabetes and fasting proinsulin adjusted for BMI were obtained from the Type 2 Diabetes / Common Metabolic Diseases Knowledge Portal (t2d.hugeamp.org/downloads). Fasting insulin adjusted for BMI summary statistics were downloaded from the MAGIC consortium ([MAGIC downloads](https://magicinvestigators.org/downloads/index.html)). Fasting plasma glucose and 2-hour post-challenge glucose summary statistics were obtained from the IEU OpenGWAS project (ebi-a-GCST90002232, ebi-a-GCST90002227). LD reference data were obtained from the publicly available UK Biobank Linkage Disequilibrium Matrices (UKBB-LD), which provide pre-computed LD matrices based on UK Biobank participants of British ancestry. TWAS prediction models are available from [PredictDB](https://predictdb.org/). The R package APCH implementing our method is available at [GitHub](https://github.com/colinyuxin/APCH).

## Notes

### Competing Interest Statement

The authors have declared no competing interest.

### Summary of Updates

Revised text, figures and supplementary materials, updated author affiliations, and released the R package.

