## Supplementary Information for "Adaptive Partial Conjunction Hypothesis: Identifying Pleiotropy Across Heterogeneous Effect Units"

Yuxin Li[1,2], Zicheng Lu[1], and Xiaolei Lin[1,*]

[1]School of Data Science, Fudan University, Shanghai, China
[2]Quantitative Biomedical Science Program, Dartmouth College, Lebanon, NH, USA

### Supplementary Contents

**1 Supplementary Notes** **3**

S1 Extended Method and Implementation Details . . . 3

S1.1 Workflow of APCH . . . 3

S1.2 PCH, APCH, and sign-aware posterior quantities . . . 3

S1.3 Gaussian approximation for summary statistics . . . 7

S1.4 Marginal model and grid distillation . . . 7

S1.5 Parameter estimation with a Dirichlet prior . . . 8

S2 Details of Simulation Studies . . . 9

S2.1 Detailed data-generating process . . . 9

S2.2 Using other methods on simulation studies . . . 10

S3 Supplementary Simulation Studies . . . 12

S3.1 Misspecification of estimation-error correlation . . . 12

S4 Details of Real Data Analysis . . . 12

S4.1 Meta-analysis of five T2D-related traits . . . 12

S4.2 Joint analysis of multi-tissue TWAS . . . 14

**2 Supplementary Tables** **17**

Table S1. GWAS summary statistics . . . 17

Table S2. APCH tag SNP counts under LD-thinning thresholds . . . 18

Table S3. Locus proxy recovery under LD-thinning thresholds . . . 18

Table S4. APCH discoveries under different SNP-selection strategies . . . 19

Table S5. Comparison of APCH-b, APCH-s, and APCH-$s_{full}$ . . . 19

Table S6. Comparison of APCH discoveries under lfsr and lfdr . . . 20

Table S7. Marginal fits and grid distillation . . . 20

Table S8. Jointly significant genes in the multi-tissue TWAS . . . 21

**3 Supplementary Figures** **22**

Figure S1. False discovery control and power in Simulation Study 1 . . . 22

Figure S2. Distribution of FDP and FSP in Simulation Study 2 . . . 23

Figure S3. Comparison of APCH-b and APCH-s in Simulation Study 3 . . . 24

Figure S4. Effect of misspecifying the estimation-error correlation on APCH . . . 25

Figure S5. Pairwise lfdr–lfsr comparison for GWAS traits . . . 26

Figure S6. Pairwise lfdr–lfsr comparison for TWAS tissues . . . 27

Figure S7. Regional association plots for APCH-selected loci . . . 28

### 1 Supplementary Notes

#### S1 Extended Method and Implementation Details

##### S1.1 Workflow of APCH

The APCH analysis proceeds through the following steps.

1. **Input data.** For each effect unit $i$, such as a SNP, gene, or locus, and each feature $j$, such as a trait, tissue, cell type, or omics measurement, APCH takes as input the marginal effect estimate $\hat{b}_{ij}$, its standard error $s_{ij}$, and an estimated or derived estimation-error correlation matrix across features. If a feature is missing for an effect unit, we treat the missing feature as missing at random and compute the likelihood using the observed features for that effect unit. APCH is intended for a moderate number of features, typically $p \leq 8$. We work with $z_{ij} = \hat{b}_{ij}/s_{ij}$ and set the marginal standard errors to one.

2. **Fit marginal model.** For each feature, we fit ash to the $z$-scores to estimate the null proportion and the non-null effect-size distribution. Before fitting the joint model, we prune and distill the variance grids. The resulting feature-wise grids provide the plug-in magnitude states for the joint APCH model.

3. **Fit joint model.** We introduce APCH-b because many existing PCH-related methods fall into this paradigm but still have limitations. Our primary model is APCH-s, which estimates the joint distribution of allocation states, where each active feature is assigned to one of its marginal ash slabs. For $p \geq 5$, we first fit second- and third-order APCH-s subproblems to prune unsupported allocation states before the final joint fit. Unless otherwise stated, APCH refers to the pruned APCH-s model.

4. **Compute subset-level posterior quantities and select discoveries.**

   From the fitted joint model, APCH computes subset-level lfdr and lfsr for each effect unit and each candidate feature subset. The lfdr is cheap to compute and is used to screen candidate subsets, so that the lfsr need not be evaluated for every subset. Level-by-level inference then selects effect units from larger to smaller candidate subsets.

##### S1.2 PCH, APCH, and sign-aware posterior quantities

**Connection to classical PCH** Classical partial conjunction hypothesis (PCH) testing evaluates whether a subset of a collection of base hypotheses is non-null. Consider $p$ base hypotheses $H_{0,1}, \ldots, H_{0,p}$, and let $r^\star = \sum_{j=1}^{p} \mathbf{1}\{H_{0,j} \text{ is false}\}$ denote the number of non-null base hypotheses. For $r \in$

$\{2, \dots, p\}$, the classical partial conjunction null is

$$H_0^{r/p} : r^\star < r.$$

$H_0^{r/p}$ states that at least $p - r + 1$ of the base null hypotheses are true, whereas its complement states that at least $r$ of the base hypotheses are false.

Liang et al. [1] noted that multiple-testing approaches to PCH can be broadly divided into empirical Bayes methods and filtering methods; see their paper for a more detailed discussion. The key idea behind EB methods for PCH is to model binary configurations. As shown below, both classical PCH-lfdr ($\mathrm{lfdr}^{\mathrm{pch}}$) and APCH subset-level lfdr ($\mathrm{lfdr}^{\mathrm{apch}}$) can be derived as linear functions of the weights of configurations. This shows that classical PCH lfdr and APCH subset-level lfdr are computed from the same type of configuration posterior, but summarize different claims. In this sense, the APCH model and other empirical Bayes methods for PCH/APCH inference follow a common modeling philosophy, with the main difference lying in how the configuration posterior is parameterized.

Recall that the binary configuration vector $\boldsymbol{\gamma}_i = (\gamma_{i1}, \dots, \gamma_{ip})$ records the null/non-null status of effect unit $i$ across the $p$ features, where $\gamma_{ij} = 1$ indicates that feature $j$ is non-null and $\gamma_{ij} = 0$ indicates that it is null. Define the posterior probability of configuration $\boldsymbol{\gamma}$ as

$$\mathrm{PPC}_i(\boldsymbol{\gamma}) = \Pr(\boldsymbol{\gamma}_i = \boldsymbol{\gamma} \mid \hat{\boldsymbol{b}}_i), \qquad \boldsymbol{\gamma} \in \{0,1\}^p.$$

Thus, the corresponding PCH-lfdr can be written as

$$\mathrm{lfdr}_i^{\mathrm{pch}}(r) = \Pr\left( \sum_{j=1}^{p} \gamma_{ij} < r \mid \hat{\boldsymbol{b}}_i \right) = \sum_{\ell=0}^{r-1} \sum_{\boldsymbol{\gamma}: \sum_{j=1}^{p} \gamma_j = \ell} \mathrm{PPC}_i(\boldsymbol{\gamma}).$$

For APCH, the subset-level lfdr asks whether every feature in this reported subset is non-null:

$$\mathrm{lfdr}_i^{\mathrm{apch}}(U) = \Pr\Big( \exists j \in U : \gamma_{ij} = 0 \mid \hat{\boldsymbol{b}}_i \Big) = \sum_{\boldsymbol{\gamma}: \exists j \in U,\ \gamma_j = 0} \mathrm{PPC}_i(\boldsymbol{\gamma}).$$

**Sign-aware inference and subset-level lfsr** A similar but more subtle connection can be made for sign-aware inference. The local false sign rate was first introduced in the ash model as the posterior probability of making a sign error [2]. For a scalar effect $b_i$, this is

$$\mathrm{lfsr}_i^{\mathrm{ash}} = \min\Big\{ \Pr(b_i \le 0 \mid \hat{b}_i), \Pr(b_i \ge 0 \mid \hat{b}_i) \Big\}.$$

Multivariate methods such as Mash, empirical Bayes matrix factorization, mvSuSiE, and mfSuSiE use the same coordinate-level sign-error idea, but condition on multivariate data [3–6]. We write this as

$$\mathrm{lfsr}_{ij}^{\mathrm{coord}} = \min\left\{\Pr(b_{ij} \le 0 \mid \hat{\boldsymbol{b}}_i), \Pr(b_{ij} \ge 0 \mid \hat{\boldsymbol{b}}_i)\right\}.$$

In this paper, consistent with the main text, we define a subset-level lfsr, denoted by $\mathrm{lfsr}^{\mathrm{apch}}$, for the claim that all features in a specified subset are non-null with a stable joint sign pattern.

Sign-aware inference has an earlier counterpart in replicability analysis (RepFDR), where the goal is to determine whether the same feature shows non-null effects with consistent directions across independent studies [7]. Heller and Yekutieli split the prior for each feature into null, positive, and negative components, which can be understood as an extension of the "three-group model" and is close in spirit to APCH-b. In this formulation, each feature is assigned to one of three states,

$$\boldsymbol{\zeta}_i = (\zeta_{i1}, \ldots, \zeta_{ip}) \in \{-1, 0, 1\}^p,$$

where $\zeta_{ij} = 1$ indicates a positive non-null effect, $\zeta_{ij} = -1$ indicates a negative non-null effect, and $\zeta_{ij} = 0$ indicates a null effect. The posterior probability of a signed configuration is

$$\mathrm{PPC}_i^{\pm}(\zeta) = \Pr(\boldsymbol{\zeta}_i = \zeta \mid \hat{\boldsymbol{b}}_i), \qquad \zeta \in \{-1, 0, 1\}^p.$$

For a signed configuration $\zeta$, let $N_+(\zeta) = \sum_{j=1}^p \mathbf{1}(\zeta_j = 1), N_-(\zeta) = \sum_{j=1}^p \mathbf{1}(\zeta_j = -1)$, and define the sign-aware PCH null set

$$\mathcal{C}_0^{\mathrm{pch}}(r) = \{\zeta \in \{-1, 0, 1\}^p : \max\{N_+(\zeta), N_-(\zeta)\} < r\}.$$

Then the sign-aware PCH error probability for the claim that at least $r$ features are non-null with a common direction can be written as

$$\mathrm{lfsr}_i^{\mathrm{pch}}(r) = \sum_{\zeta \in \mathcal{C}_0^{\mathrm{pch}}(r)} \mathrm{PPC}_i^{\pm}(\zeta).$$

For a signed configuration $\zeta$, let $\zeta_U$ denote its restriction to the coordinates in $U$. Then

$$\mathrm{lfsr}_i^{\mathrm{apch}}(U) = 1 - \max_{s \in \{\pm 1\}^{|U|}} \sum_{\zeta : \zeta_U = s} \mathrm{PPC}_i^{\pm}(\zeta).$$

The signed-configuration notation above is most direct when the prior itself contains separate null, positive, and negative components. This separation is not required. Suppose the model first defines a binary null/non-null configuration $\boldsymbol{\gamma}_i \in \{0, 1\}^p$. For a signed configuration $\zeta \in \{-1, 0, 1\}^p$, let $S_\zeta = \{j : \zeta_j \neq 0\}$ be the set of active coordinates encoded by $\zeta$. Given $\boldsymbol{\gamma}_i = \mathbf{1}_{S_\zeta}$, suppose the active effects $\boldsymbol{b}_{i,S_\zeta}$ have a proper posterior density $p_i\Big(\boldsymbol{b}_{S_\zeta} \mid \hat{\boldsymbol{b}}_i, \boldsymbol{\gamma}_i = \mathbf{1}_{S_\zeta}\Big)$ on $\mathbb{R}^{|S_\zeta|}$. Then

$$\mathrm{PPC}_i^{\pm}(\zeta) = \Pr\Big(\boldsymbol{\gamma}_i = \mathbf{1}_{S_\zeta} \mid \hat{\boldsymbol{b}}_i\Big) \int_{\{\boldsymbol{b}_{S_\zeta}:\, \zeta_j b_{ij} > 0,\ j \in S_\zeta\}} p_i\Big(\boldsymbol{b}_{S_\zeta} \mid \hat{\boldsymbol{b}}_i, \boldsymbol{\gamma}_i = \mathbf{1}_{S_\zeta}\Big)\ d\boldsymbol{b}_{S_\zeta}.$$

Thus the positive and negative signs can be introduced by integrating the conditional posterior distribution over the corresponding orthant.

If a model has no explicit null/non-null configuration, the same formula can be viewed as the special case $\boldsymbol{\gamma}_i = (1, \ldots, 1)$. Then $\mathrm{PPC}_i^{\pm}(\zeta)$ is still well defined. However, the sign-aware PCH quantity can become trivial. For example, when $p = 3$ and $r = 2$, every vector in $\{-1, 1\}^3$ has at least two coordinates with the same sign. Hence $\mathrm{lfsr}_i^{\mathrm{pch}}(2) = 0$ regardless of the data.

This construction also clarifies the relationship between $\mathrm{lfsr}^{\mathrm{apch}}$ and coordinate-level lfsr. When $U = \{j\}$,

$$\mathrm{lfsr}_i^{\mathrm{apch}}(\{j\}) = 1 - \max_{s \in \{-1,1\}} \Pr(\zeta_{ij} = s \mid \hat{\boldsymbol{b}}_i).$$

Since

$$\Pr(\zeta_{ij} = 1 \mid \hat{\boldsymbol{b}}_i) = \Pr(b_{ij} > 0 \mid \hat{\boldsymbol{b}}_i), \qquad \Pr(\zeta_{ij} = -1 \mid \hat{\boldsymbol{b}}_i) = \Pr(b_{ij} < 0 \mid \hat{\boldsymbol{b}}_i),$$

we have

$$\begin{aligned}\mathrm{lfsr}_i^{\mathrm{apch}}(\{j\}) &= 1 - \max\Big\{\Pr(b_{ij} > 0 \mid \hat{\boldsymbol{b}}_i), \Pr(b_{ij} < 0 \mid \hat{\boldsymbol{b}}_i)\Big\} \\ &= \min\Big\{\Pr(b_{ij} \le 0 \mid \hat{\boldsymbol{b}}_i), \Pr(b_{ij} \ge 0 \mid \hat{\boldsymbol{b}}_i)\Big\} = \mathrm{lfsr}_{ij}^{\mathrm{coord}}.\end{aligned}$$

The derivation above shows that $\mathrm{lfsr}^{\mathrm{apch}}$ is a flexible posterior quantity and is not tied to a specific prior formulation. The three computational forms above cover the main cases: (1) an explicit signed prior; (2) a prior with a null component and a continuous non-null component; and (3) a fully continuous prior.

RepFDR with signed configurations falls into case (1), whereas APCH belongs to case (2); fully continuous multivariate shrinkage models, such as Mash, fall closer to case (3).

We advocate subset-level lfsr as a more informative posterior quantity than coordinate-level lfsr in multivariate settings. A small $\mathrm{lfsr}^{\mathrm{apch}}$ indicates strong posterior evidence that the features in the reported subset have a stable joint sign pattern. However, $\mathrm{lfsr}^{\mathrm{apch}}$ alone does not guarantee calibrated error-rate control. Although a fully continuous prior can still produce $\mathrm{lfsr}^{\mathrm{apch}}$, Simulation Study 2 shows that Mash can produce many false positive subset claims. Recent work also shows that coordinate-level lfsr in Mash-type multivariate empirical Bayes models can be anti-conservative [8, 9]. This suggests that an explicit binary configuration layer, with null or inactive components, provides an important form of shrinkage toward zero; without such components, a fully continuous prior may not reliably separate null and non-null effects from the data.

It remains an open and interesting question whether $\mathrm{lfsr}^{\mathrm{apch}}$ computed from a binary null/non-null prior with continuous non-null effects can provide calibrated false sign control. We therefore view

$\mathrm{lfsr}^{\mathrm{apch}}$ primarily as a robust posterior ranking quantity that shrinks weak and sign-unstable effects. Because $\mathrm{lfsr}_i^{\mathrm{apch}}(U) \geq \mathrm{lfdr}_i^{\mathrm{apch}}(U)$, selection by lfsr is never less conservative than selection by lfdr. Further work on calibrated subset-level lfdr and lfsr remains important.

##### S1.3 Gaussian approximation for summary statistics

APCH assumes the observed marginal test statistics are normally distributed around the true value and is therefore most applicable to quantitative traits. Provided the effect sizes are small, there is good theoretical and empirical justification for applying standard linear models directly to binary traits [10, 11]. The Gaussian approximation is therefore also appropriate for binary traits. If the sample size is small, the noise can be better modeled as a $t$-distribution [2]. However, in current association studies, the sample size is usually moderate to large. More generally, this approximation means that APCH is most suitable to deal with data that are generated feature by feature from linear regression models. Still, there has been little theoretical or empirical assessment to justify this approximation when the summary statistics are generated from more complex models such as linear mixed models, generalized linear models, or even non-linear models.

The estimation-error correlation matrix $\boldsymbol{C}_i$ describes the dependence between residual errors. This correlation can be substantially large in our multi-tissue TWAS application, where the test statistics of different tissues for the same gene are computed from overlapping SNPs. Since $\boldsymbol{C}_i$ is a plug-in estimator and inference can be sensitive to plug-in error in $\boldsymbol{C}_i$ when it is near singular (see the derivation and simulation below), we place a floor of $0.05$ on the smallest eigenvalue of $\boldsymbol{C}_i$:

$$\widetilde{\boldsymbol{C}}_i = (1-\alpha_i)\boldsymbol{C}_i + \alpha_i \boldsymbol{I}, \qquad \alpha_i = \max\left\{0,\, \frac{0.05 - \lambda_{\min}(\boldsymbol{C}_i)}{1-\lambda_{\min}(\boldsymbol{C}_i)}\right\}.$$

A near-singular $\boldsymbol{C}_i$ can make the Gaussian likelihood overly sensitive to small contrast directions. For any unit vector $v$, the error variance of the contrast $v^\top z_i$ is $v^\top \boldsymbol{C}_i v$. Hence a small $\lambda_{\min}(\boldsymbol{C}_i)$ means that some contrast is treated as nearly noise-free. For example, suppose two features have error correlation $\rho = 0.999$ and we observe $z = (0.1, -0.1)$. Although both marginal $z$-scores are small, the contrast is $z_1 - z_2 = 0.2$, while its error variance is only $2(1-\rho) = 0.002$. So the plug-in Gaussian likelihood treats this small marginal pattern as evidence for $b_1 - b_2 \neq 0$.

##### S1.4 Marginal model and grid distillation

**Marginal grid distillation** For each feature $j$, we fit ash [2] on a dense geometric grid $\sigma_{\min,j} \leq \sigma_{j1} < \cdots < \sigma_{jK_j} \leq \sigma_{\max,j}$, where $\sigma_{\min,j} = 0.1\,\mathrm{median}_i(s_{ij})$ and

$$\sigma_{\max,j} = \begin{cases} 2\left\{\max_i\left((\hat{b}_{ij})^2 - s_{ij}^2\right)\right\}^{1/2}, & \text{if } \max_i\left((\hat{b}_{ij})^2 - s_{ij}^2\right) > 0, \\ 12\,\sigma_{\min,j}, & \text{otherwise.} \end{cases}$$

We then remove nonzero-variance components with negligible conditional non-null weight $q_{jk} < 10^{-6}$ and further compress the non-null grid by iterative merging, following the general idea of KL-based Gaussian mixture reduction [12]. Let $v_{jk} = \sigma^2_{jk}$ and sort the nonzero components by $v_{jk}$. For adjacent pairs $(k, k+1)$, we define the merge cost as the weighted symmetric KL divergence between the corresponding normal components and prioritize pairs with the smallest cost. Merging replaces $(v_{jk}, v_{j,k+1})$ by $v_{\text{new}} = (q_{jk}v_{jk} + q_{j,k+1}v_{j,k+1})/(q_{jk} + q_{j,k+1})$ while leaving the null component unchanged. At each iteration, we try adjacent merges in order of increasing cost, refit all mixture weights on the reduced grid, and accept the first merge that decreases the mean log-likelihood per effect by less than $2 \times 10^{-4}$.

##### S1.5 Parameter estimation with a Dirichlet prior

The Dirichlet prior used in APCH is intended as weak pseudo-count regularization for finite-mixture weights. We consider a symmetric Dirichlet prior $w \sim \text{Dir}(\alpha, \ldots, \alpha)$ with $\alpha > 1$. For $\alpha$ slightly above 1, little estimation bias is induced, while the variance decreases.

Let $w = (w_1, \ldots, w_m)$ be the mixture weight vector, and write $w = w^\star + B\theta$ locally around the truth $w^\star$. Around the MLE $\hat{\theta}_{\text{MLE}}$, suppose the negative log-likelihood is locally quadratic with Hessian $H_n$. Adding pseudo-counts contributes an additional positive-curvature term, so the regularized solution has the local form

$$\hat{\theta}_{\text{reg}} \approx (H_n + P_n)^{-1} H_n \hat{\theta}_{\text{MLE}},$$

where $P_n$ is positive semidefinite. Thus, symmetric pseudo-count regularization acts locally like ridge shrinkage: for pseudo-count strength $\lambda$, the squared bias is $O(\lambda^2)$, whereas the variance is reduced at first order in $\lambda$. Therefore, for sufficiently small pseudo-count strength, the overall mean squared error can be smaller than that of the MLE.

The pseudo-count for an allocation state $\boldsymbol{h}$ is set to

$$\alpha_{\boldsymbol{h}} = \max\left\{10^{-4},\, \min(0.001\, n\, q_{\boldsymbol{h}},\, 5)\right\}, \qquad q_{\boldsymbol{h}} \propto \prod_{j:h_j=0} p_{0j} \prod_{j:h_j>0} (1 - p_{0j})\hat{q}_{jh_j}.$$

Here $p_{0j}$ is the fitted marginal null probability and $\hat{q}_{jk}$ is the estimated feature-wise conditional mixture weight from the marginal fit. For allocation-state pruning in APCH-s, we fit pairwise and three-way APCH-s submodels. A local allocation state $\boldsymbol{h}$ is pruned if its effective posterior count

$$\widehat{N}(\boldsymbol{h}) = \sum_{i=1}^{n} r_{i\boldsymbol{h}}, \qquad r_{i\boldsymbol{h}} = \Pr(\boldsymbol{\eta}_i = \boldsymbol{h} \mid \hat{\boldsymbol{b}}_i),$$

satisfies

$$\widehat{N}(\boldsymbol{h}) < 1 \quad \text{and} \quad \widehat{N}(\boldsymbol{h})/\alpha_{\boldsymbol{h}} < 10.$$

Every full allocation state compatible with a pruned low-order state is then removed from the final APCH-s support.

### S2 Details of Simulation Studies

#### S2.1 Detailed data-generating process

**Simulation Study 1** We generated $n = 10{,}000$ independent effects with $p = 2$ traits. Each effect first drew a binary configuration $\boldsymbol{\gamma}_i \in \{00, 10, 01, 11\}$. In the mixed configuration design, the probabilities for $00, 10, 01, 11$ were $0.80, 0.10, 0.05, 0.05$. In the restricted design, only $00$ and $11$ were allowed, with $\varphi_{00} \in \{0.90, 0.95, 0.99, 0.995\}$.

Given $\boldsymbol{\gamma}_i$, inactive coordinates were set to zero and active coordinates were sampled independently from one of three marginal effect-size distributions. The normal setting used $N(0, 1)$. The near-normal setting used $\frac{2}{3}N(0, 0.75^2) + \frac{1}{3}N(0, 1.5^2)$. The spiky setting used $0.2N(0, 0.25^2) + 0.4N(0, 0.5^2) + 0.2N(0, 1) + 0.2N(0, 2^2)$.

Observed effects were generated as $\hat{\boldsymbol{b}}_i = \boldsymbol{b}_i + \boldsymbol{\varepsilon}_i$, where $\boldsymbol{\varepsilon}_i \sim N_2(0, 0.35^2 R_\rho)$, $R_\rho = (1-\rho)I_2 + \rho\mathbf{1}\mathbf{1}^\top$. We used $\rho = 0.9$ for the normal setting, $\rho = 0.3$ for the near-normal setting, and $\rho = 0$ for the spiky setting.

**Simulation Study 2** We generated $n = 100{,}000$ independent effects with $p = 5$ traits. The true effects were generated from structural factor models.

In Scenario 1, two latent factors generated the effects: $\boldsymbol{b}_i = A_{i1}(1, 1, 1.5, 0, 0)^\top + A_{i2}(0, 0, 0.5, 2, 1)^\top$, where $A_{i1} \sim N(0, 6)$ and $A_{i2} \sim N(0, 4)$. The factor on/off counts for $00, 10, 01, 11$ were $95{,}000, 2{,}000, 2{,}000, 1{,}000$.

In Scenario 2, the same two-factor structure was used, and trait-specific marginal signals were added. For each trait, $5{,}000$ disjoint effects were assigned a single-trait effect sampled from $N(0, 1.5)$, with all other coordinates set to zero.

In Scenario 3, three overlapping latent factors generated the structured effects:
$\boldsymbol{b}_i = A_{i1}(1.2, 1, 1.1, 0.9, 0)^\top + A_{i2}(0, 1.1, 0.9, 1, 1.2)^\top + A_{i3}(1, 1.2, 0, 1, 0.9)^\top$, where $A_{i1} \sim N(0, 6)$, $A_{i2} \sim N(0, 4)$, and $A_{i3} \sim N(0, 3.5)$. The factor on/off counts for $000, 100, 010, 001, 110, 101, 011, 111$ were $95{,}000, 1{,}100, 1{,}100, 1{,}100, 400, 400, 400, 500$. The same trait-specific marginal signals as in Scenario 2 were then added.

In all three scenarios, observed effects were generated as $\hat{\boldsymbol{b}}_i = \boldsymbol{b}_i + \boldsymbol{\varepsilon}_i$, where $\boldsymbol{\varepsilon}_i \sim N_5(0, I_5)$.

**Simulation Study 3** We generated $n = 100{,}000$ independent effects with $p = 2$ traits. The configuration counts for $00, 10, 01, 11$ were fixed at $98{,}500, 600, 600, 300$. Inactive coordinates were set to zero. Active effects were generated from three Gaussian slabs with scales $\tau = (0.5, 1, 2)$, with equal slab probabilities across all scenarios. The marginal-only configurations $10$ and $01$ were kept the same across all scenarios: conditional on the configuration, the single active coordinate was sampled from its assigned slab and the other coordinate was set to zero.

The four scenarios differed only in the joint non-null configuration 11. In Scenario 1, conditional on $\gamma_i = 11$, the two active coordinates had independent allocation states: $b_{ij} \mid k_{ij} \sim N(0, \tau^2_{k_{ij}})$ for $j = 1, 2$, with $k_{ij}$ sampled uniformly from $\{1, 2, 3\}$.

In Scenario 2, the two active coordinates shared the same allocation state $\eta_i = (k_i, k_i)$, with $k_i$ sampled uniformly from $\{1, 2, 3\}$. Conditional on this allocation vector, the effect values were independent, with $b_{ij} \mid k_i \sim N(0, \tau^2_{k_i})$ for $j = 1, 2$.

In Scenario 3, the two active coordinates again shared the same allocation state $\eta_i = (k_i, k_i)$, with $k_i$ sampled uniformly from $\{1, 2, 3\}$, but the effect values were additionally correlated: $\boldsymbol{b}_i \mid k_i \sim N_2(0, \tau^2_{k_i} R_b)$, where $R_b$ has unit diagonal and off-diagonal correlation $\rho_b = 0.5$.

In Scenario 4, the two active coordinates shared the same allocation state, with $k_i$ sampled uniformly from $\{1, 2, 3\}$, and were perfectly correlated. We sampled $a_i \sim N(0, \tau^2_{k_i})$ and set $\boldsymbol{b}_i = (a_i, a_i)$. This is the $\rho_b = 1$ limit of Scenario 3.

In all four scenarios, observed effects were generated as $\hat{\boldsymbol{b}}_i = \boldsymbol{b}_i + \boldsymbol{\varepsilon}_i$, where $\boldsymbol{\varepsilon}_i \sim N_2(0, 0.25^2 I_2)$.

##### S2.2 Using other methods on simulation studies

RepFDR [7] and OPERA [13] are the clearest examples of the APCH-b regime, although they use different priors for the marginal effect-size distributions. Furthermore, RepFDR, to our knowledge, was the first to put forward the idea of an "extension of the two-group model" by modeling the binary configuration, although it cannot explicitly model the estimation-error correlation. CPBayes [14] is a Bayesian subset-based method that also models trait-specific association configurations and estimates their posterior probabilities. The non-null effects across traits are modeled with a shared slab, so it is also closer in spirit to APCH-b. PLACO [15] gave a clear formulation of why PCH is important for pleiotropy detection by directly targeting joint non-null evidence under a composite null, while PLACO+ [16] extends this idea in a more robust way to account for estimation-error correlation. However, both methods are limited to two traits, and their sub-null distributions are estimated by pooling feature-wise data only. We also note that earlier methods such as ASSET [17] and GPA [18] are conceptually related, especially ASSET, which is an important early subset-based approach closely connected to the PCH objective. These earlier methods have already been discussed extensively in the CPBayes literature, so we do not include them in the main comparisons.

Mash [3], and its improved implementation UDR [8], are closely related to APCH. Mash systematically developed the empirical Bayes multivariate normal means framework for multivariate effect estimation and testing, and UDR provides an improved implementation of this framework. They model the joint effect-size distribution through the prior

$$\boldsymbol{b}_i \sim \sum_{\ell=1}^{L} \pi_\ell \mathcal{N}_p(\mathbf{0}, \boldsymbol{U}_\ell), \qquad \hat{\boldsymbol{b}}_i \mid \boldsymbol{b}_i \sim \mathcal{N}_p(\boldsymbol{b}_i, \boldsymbol{E}_i).$$

These methods are highly effective at learning high-dimensional sharing patterns through the covariance matrices $\boldsymbol{U}_\ell$, but they do not explicitly parameterize the $2^p$ binary significance configurations that are central to the PCH objective. In this paper, we evaluate Mash/UDR only in simulations with independent noise. To assess subset-level joint evidence, we compute a subset-level lfsr from the fitted mixture posterior. Let

$$\xi_{i\ell} = \frac{\pi_\ell\, \varphi_p(\hat{\boldsymbol{b}}_i; \mathbf{0}, \boldsymbol{U}_\ell + \boldsymbol{E}_i)}{\sum_{m=1}^{L} \pi_m\, \varphi_p(\hat{\boldsymbol{b}}_i; \mathbf{0}, \boldsymbol{U}_m + \boldsymbol{E}_i)}$$

denote the posterior responsibility of component $\ell$ for effect $i$, and let $\boldsymbol{b}_i \mid \hat{\boldsymbol{b}}_i, \ell \sim \mathcal{N}_p(\boldsymbol{\mu}_{i\ell}, \boldsymbol{\Sigma}_{i\ell})$ be the corresponding Gaussian posterior, with

$$\boldsymbol{\mu}_{i\ell} = \boldsymbol{U}_\ell(\boldsymbol{U}_\ell + \boldsymbol{E}_i)^{-1}\hat{\boldsymbol{b}}_i, \qquad \boldsymbol{\Sigma}_{i\ell} = \boldsymbol{U}_\ell - \boldsymbol{U}_\ell(\boldsymbol{U}_\ell + \boldsymbol{E}_i)^{-1}\boldsymbol{U}_\ell.$$

Writing $\mathcal{A}_U = \{\ell : (\boldsymbol{U}_\ell)_{jj} > 0 \text{ for all } j \in U\}$ and $\mathcal{B}(U, s) = \{\boldsymbol{b} :\ s_j b_j > 0 \text{ for all } j \in U\}$, we define

$$\mathrm{lfsr}_i^{\mathrm{mash}}(U) = 1 - \max_{s \in \{\pm 1\}^{|U|}} \sum_{\ell \in \mathcal{A}_U} \xi_{i\ell}\, \Pr\big(\boldsymbol{b}_{i,U} \in \mathcal{B}(U, s) \mid \hat{\boldsymbol{b}}_i, \ell\big).$$

For CPBayes in the two-trait simulations, we use its posterior distribution on the binary configuration $\gamma_i \in \{0, 1\}^2$ to derive the corresponding subset-level lfdrs. We apply BH adjustment to PLACO p-values, and use the subset-level lfdrs from OPERA and RepFDR in the same level-by-level inference for fair FDR comparison.

It is also noteworthy that none of these methods, whether they learn the effect-size distribution from the data or test effect units one by one, can directly model or adjust for correlation between effect units, such as that induced by LD.

We ran RepFDR, binning z-scores with ztobins using two states and obtaining per-configuration posteriors with ldr. We increased the degrees of freedom from 35 to 84 in steps of 7 and chose the smallest $df$ with misfit $\leq 1$; if none satisfied this rule, we used $df = 84$. We ran OPERA under default settings. We ran PLACO, estimating the marginal variance with var.placo; in correlated simulations we used placo.plus, whereas in uncorrelated settings we used placo. We ran CPBayes with the correlated and uncorrelated models through cpbayes_cor and cpbayes_uncor, respectively. Each analysis used 8,000 total MCMC iterations and a burn-in of 2,000 iterations. In simulations with correlated noise, we supplied the true correlation matrix for each method. We ran mashr and its improved implementation udr. Following the improved UDR pipeline, we used truncated eigenvalue decomposition (TED) updates, inverse-Wishart regularization with $\lambda = p$, and a random initialization consisting of a fixed global-null covariance matrix and randomly initialized unconstrained covariance components. For the three scenarios in Simulation Study 2, we used $1+10$, $1+15$, and $1+20$ mixture components, respectively.

### S3 Supplementary Simulation Studies

#### S3.1 Misspecification of estimation-error correlation

**Experimental setup** To examine sensitivity to plug-in error in the estimation-error correlation, we simulated 2 features with a fixed APCH prior. Supplementary Figure 4 summarizes the results. Let $\boldsymbol{\gamma}_i \in \{00, 10, 01, 11\}$ denote the binary configuration, with $(\pi_{00}, \pi_{10}, \pi_{01}, \pi_{11}) = (0.80, 0.08, 0.08, 0.04)$. Given $\boldsymbol{\gamma}_i = \boldsymbol{h}$, we used $\boldsymbol{b}_i \mid \boldsymbol{\gamma}_i = \boldsymbol{h} \sim N_2(0, D_{\boldsymbol{h}})$ with $D_{\boldsymbol{h}} = \mathrm{diag}(4h_1, 4h_2)$. We considered true correlations $\rho \in \{0.90, 0.95, 0.99, 0.9999\}$ and evaluated posterior quantities using plug-in correlations $\hat{\rho} \in \{0, 0.90, 0.95, 0.99, 0.9999\}$. Panels A–C use this fixed prior to generate samples and compute posterior quantities. For Panel D, we refitted the configuration proportions with the slab variance fixed as known parameters.

**Results** Under strong estimation-error correlation, mild underestimation of the correlation is safer than overestimation (Supplementary Figure 4A, B, and D). We therefore place a floor of $0.05$ on the smallest eigenvalue of $\boldsymbol{C}_i$ in the primary analysis. At the same time, the estimation-error correlation contains useful information for the EB model, especially when the true correlation is large, because it encodes the directions in which noise is expected to vary (Supplementary Figure 4C).

Treating strongly correlated errors as independent is also problematic. First, it is not uniformly conservative: when the correctly specified posterior probability of the global-null configuration is large, using $\hat{\rho} = 0$ tends to underestimate this probability. Second, it can also distort prior-weight estimation.

We also considered the case of negative error correlations and obtained consistent results.

### S4 Details of Real Data Analysis

#### S4.1 Meta-analysis of five T2D-related traits

We analyzed GWAS summary statistics for five T2D-related traits in individuals of European ancestry (Supplementary Table 1).

**Data processing** We first restricted to autosomal, biallelic SNPs and removed strand-ambiguous (A/T or C/G) and multi-allelic variants, SNPs with minor allele frequency (MAF) $< 0.01$ in any study, and variants in the extended MHC region. We then harmonized alleles across traits by aligning all effect and non-effect alleles to the T2D GWAS, flipping effect signs when necessary and discarding SNPs with inconsistent or unresolvable allele labels. We further restricted to SNPs that were observed (non-missing) in all five traits.

**Estimating the correlation matrix** The estimation-error correlation mainly reflects sample overlap that is shared across SNPs, so we assume a common correlation structure and set $\boldsymbol{C}_i = \boldsymbol{C}$ for all SNPs. We started from SNPs that were non-missing in all five traits and estimated $\boldsymbol{C}$ using near-independent null variants. After LD clumping the T2D GWAS at $r^2 < 0.01$, we selected 25,845 SNPs with $p$-values greater than $0.1$ in all five traits and computed the sample correlation matrix of their effect estimates. This revealed a modest correlation only between FI and FPG ($0.1347$), with all other pairwise correlations below $0.05$. Repeating the same procedure with a more permissive clumping threshold of $r^2 < 0.25$ again gave correlations below $0.05$ for all trait pairs other than FI–FPG ($0.1347$), but these small estimates varied noticeably, suggesting that they largely reflect estimation noise. We therefore retained the FI–FPG correlation and set all other off-diagonal elements of $\boldsymbol{C}$ to zero.

**Sensitivity analysis of different SNP-selection strategies** We conducted a sensitivity analysis to assess how APCH depends on the choice of LD-thinning threshold and SNP-selection strategy. First, under random thinning, we compared the numbers of $L = 3$, $L = 4$, and $L = 5$ tag SNP discoveries across $r^2 = 0.10$, $0.25$, and $0.60$ using subset-level lfsr at $\alpha = 0.01$ (Supplementary Table 2).

Second, to examine whether wider thinning mainly recovers new loci or simply retains more correlated proxies, we focused on five well-established T2D loci with relatively clear literature-supported sentinel or functional variants: *SLC30A8* rs13266634, *TCF7L2* rs7903146, *CDKAL1* rs7756992, *ADCY5* rs11708067, and *GLIS3* rs7034200 [19–22]. For *GLIS3*, rs7034200 was not present in the UKBB LD panel used here. We therefore used rs3892354 as a proxy anchor, which is in very high LD with rs7034200 ($r^2 \approx 0.97$). For each locus, we counted APCH discoveries under lfsr-based inference at $\alpha = 0.05$ that were in LD ($r^2 \geq 0.25$) with the locus anchor (Supplementary Table 3).

Third, we compared random thinning with alternative SNP-selection strategies based on single-trait lead SNPs and MetaUSAT lead SNPs (Supplementary Table 4). For a single-trait lead panel, the representative SNP in each LD region was chosen as the SNP with the smallest $p$-value for the corresponding trait. For the MetaUSAT lead panel, representative SNPs were selected using $p$-values from MetaUSAT, implemented with the software available at `https://github.com/RayDebashree/metaUSAT`. MetaUSAT is a unified score-based omnibus association test that combines evidence across multiple traits using only GWAS summary statistics.

In short, random thinning at $r^2 = 0.25$ gives a practical balance: it retains representative signals at each locus while limiting proxy multiplicity and avoids the selection bias introduced by lead-SNP-based selection.

**Comparison of joint models** We compared the performance of APCH-b, APCH-s, and APCH-s$_{\text{full}}$. APCH-s$_{\text{full}}$ refers to the APCH-s model without pruning allocation states. Supplementary Tables 5 and 6 summarize the results. Allocation-state pruning typically reduces the number of parameters from 432 to about 150 and reduces runtime by more than 20-fold while discovering almost the same SNPs compared with APCH-s$_{\text{full}}$.

Both the number and quality of APCH-b discoveries are very limited compared with APCH-s, suggesting that the joint-significance discoveries are mainly driven by large slabs. Notably, APCH-b is highly conservative under lfsr-based inference. This is because, in GWAS data, the ash model usually identifies very small slabs, for example with $\sigma^2 < 1$, and concentrates substantial mass on them. This also makes exact-zero and near-zero effects difficult to distinguish, which further motivates our use of lfsr as the primary inferential quantity.

##### S4.2 Joint analysis of multi-tissue TWAS

We followed the MetaXcan tutorials to harmonize and impute the T2D GWAS summary statistics and to compute single-tissue $z$-scores with S-PrediXcan. Following Liang et al. [23], the $z$-score of gene $i$ in tissue $j$ was calibrated as

$$\tilde{z}_{ij} = \frac{z_{ij}}{\sqrt{1 + \phi_{ij}\, Nh^2}},$$

where $\phi_{ij}$ is the gene- and tissue-specific variance parameter of the prediction model, $N = 293{,}100$ is the effective sample size of the T2D GWAS, and $h^2 = 0.15$ is the observed-scale SNP heritability estimated by LD score regression [24] (Supplementary Table 1). Let $\boldsymbol{w}_{ij}$ denote the SNP weights of the prediction model for gene $i$ in tissue $j$ and $\boldsymbol{R}$ the SNP correlation matrix of the reference panel. The estimation-error correlation between tissues $j$ and $j'$ is the correlation of the predicted expression levels,

$$(\boldsymbol{C}_i)_{jj'} = \frac{\boldsymbol{w}_{ij}^\top \boldsymbol{R}\, \boldsymbol{w}_{ij'}}{\sqrt{\boldsymbol{w}_{ij}^\top \boldsymbol{R}\, \boldsymbol{w}_{ij}}\, \sqrt{\boldsymbol{w}_{ij'}^\top \boldsymbol{R}\, \boldsymbol{w}_{ij'}}},$$

as computed by S-MultiXcan [25]. Since the calibration rescales each $z$-score by a positive constant, it leaves $\boldsymbol{C}_i$ unchanged. Genes that are not predictable in a tissue were treated as missing completely at random, and their likelihood was computed from the observed tissues only.

#### 2 Supplementary Tables

| GWAS trait | Description | Source portal | Sample size |
|---|---|---|---|
| T2D | Type 2 diabetes | `T2DKP` | 80,154 cases and 853,816 controls |
| FI | Fasting insulin adjusted for BMI | `MAGIC` | Up to 151,013 individuals |
| PROI | Fasting proinsulin adjusted for BMI | `T2DKP` | Up to 45,826 individuals |
| FPG | Fasting plasma glucose | `IEU OpenGWAS` | Up to 200,622 individuals |
| 2hPG | Two-hour post-challenge glucose | `IEU OpenGWAS` | Up to 63,396 individuals |
| Counts by number of traits with $p$-value $< 10^{-4}$ | | | |
| Number of SNPs in the primary analysis | | | 192,755 |
| Exactly 1 trait | | | 1,226 |
| Exactly 2 traits | | | 162 |
| Exactly 3 traits | | | 41 |
| Exactly 4 traits | | | 9 |
| Exactly 5 traits | | | 1 |
| TWAS: counts by number of tissues with $p$-value $< 10^{-4}$ | | | |
| Number of genes predictable in at least one tissue | | | 14,501 |
| Exactly 1 tissue | | | 60 |
| Exactly 2 tissues | | | 10 |
| Exactly 3 tissues | | | 8 |
| Exactly 4 tissues | | | 2 |
| Exactly 5 tissues | | | 3 |
| Exactly 6 tissues | | | 3 |

Supplementary Table 1: GWAS summary statistics used and counts of variants with $p$-value $< 10^{-4}$ in 1–5 traits in the primary analysis. IEU OpenGWAS IDs are `ebi-a-GCST90002232` for FPG and `ebi-a-GCST90002227` for 2hPG. TWAS prediction models are available from `PredictDB`.

| LD threshold | $L = 3$ | $L = 4$ | $L = 5$ |
|---|---|---|---|
| $r^2 = 0.10$ | 35.7 ± 4.7 (13.0%) | 15.0 ± 3.4 (22.8%) | 5.2 ± 2.1 (39.6%) |
| $r^2 = 0.25$ | 48.6 ± 5.4 (11.2%) | 19.5 ± 3.5 (18.1%) | 8.4 ± 1.9 (22.6%) |
| $r^2 = 0.60$ | 67.9 ± 5.0 (7.3%) | 25.6 ± 2.4 (9.3%) | 11.6 ± 1.4 (11.6%) |

Supplementary Table 2: APCH tag SNP counts under random LD-thinning thresholds. Results are summarized over 50 random-thinning replicates using subset-level lfsr at $\alpha = 0.01$. Entries are reported as mean ± standard deviation, with the coefficient of variation shown in parentheses, where $\mathrm{CV} = \mathrm{SD}/\mathrm{mean}$. Tag SNPs were defined as in the main text: within each LD block, we chose the SNP with (i) the largest number of traits for which APCH declared joint significance and (ii) the smallest corresponding lfsr within that block.

| Locus | $r^2 = 0.10$ | | | $r^2 = 0.25$ | | | $r^2 = 0.60$ | | |
|---|---|---|---|---|---|---|---|---|---|
| | Proxy Count | Hit | Mean $r^2$ | Proxy Count | Hit | Mean $r^2$ | Proxy Count | Hit | Mean $r^2$ |
| *SLC30A8* | 0.62 ± 0.49 | 62% | 0.812 | 1.08 ± 0.27 | 100% | 0.740 | 2.72 ± 0.61 | 100% | 0.598 |
| *TCF7L2* | 0.68 ± 0.47 | 68% | 0.573 | 1.10 ± 0.30 | 100% | 0.625 | 2.42 ± 0.50 | 100% | 0.605 |
| *CDKAL1* | 0.92 ± 0.60 | 78% | 0.451 | 2.26 ± 0.94 | 100% | 0.420 | 6.64 ± 0.72 | 100% | 0.482 |
| *ADCY5* | 0.82 ± 0.39 | 82% | 0.716 | 1.36 ± 0.48 | 100% | 0.573 | 3.00 ± 0.00 | 100% | 0.603 |
| *GLIS3* | 0.64 ± 0.48 | 64% | 0.807 | 1.00 ± 0.00 | 100% | 0.819 | 1.28 ± 0.45 | 100% | 0.747 |

Supplementary Table 3: Known-locus proxy recovery under random LD-thinning thresholds. For each established T2D locus, proxy SNPs were defined as APCH-reported SNPs using lfsr at $\alpha = 0.05$ that were in pairwise LD $r^2 \geq 0.25$ with the locus anchor. Under each thinning threshold, we report the mean ± standard deviation of the number of selected proxies across 50 random-thinning replicates, the hit rate, defined as the percentage of replicates with at least one proxy selected, and the mean $r^2$ of the retained proxies.

$r^2 = 0.10$

| | PROI | FI | T2D | FPG | 2hPG |
|---|---|---|---|---|---|
| Random thinning | 71.1 | 72.2 | 475.3 | 208.3 | 51.8 |
| PROI lead | **935** | 86 | 420 | 226 | 60 |
| FI lead | 71 | **1842** | 416 | 202 | 60 |
| T2D lead | 67 | 115 | **6374** | 277 | 67 |
| FPG lead | 105 | 85 | 459 | **2261** | 89 |
| 2hPG lead | 77 | 90 | 447 | 223 | **1017** |
| MetaUSAT lead | 138↑ | 292↑ | 1828↑ | 499↑ | 141↑ |

$r^2 = 0.25$

| | PROI | FI | T2D | FPG | 2hPG |
|---|---|---|---|---|---|
| Random thinning | 134.8 | 140.4 | 948.0 | 397.3 | 96.6 |
| PROI lead | **685** | 148 | 879 | 395 | 107 |
| FI lead | 148 | **1155** | 854 | 392 | 119 |
| T2D lead | 149 | 193 | **6205** | 477 | 132 |
| FPG lead | 157 | 153 | 915 | **1657** | 100 |
| 2hPG lead | 145 | 150 | 896 | 412 | **622** |
| MetaUSAT lead | 232↑ | 336↑ | 2052↑ | 662↑ | 120↑ |

$r^2 = 0.60$

| | PROI | FI | T2D | FPG | 2hPG |
|---|---|---|---|---|---|
| Random thinning | 304.5 | 391.3 | 2565.3 | 951.6 | 228.1 |
| PROI lead | **441** | 407 | 2609 | 987 | 222 |
| FI lead | 315 | **688** | 2579 | 979 | 246 |
| T2D lead | 323 | 409 | **6143** | 1018 | 227 |
| FPG lead | 342 | 406 | 2489 | **1301** | 140 |
| 2hPG lead | 337 | 410 | 2523 | 881 | **319** |
| MetaUSAT lead | 391↑ | 584↑ | 3534↑ | 1139↑ | 254↑ |

Supplementary Table 4: Trait-wise discovery counts from APCH using lfsr at $\alpha = 0.01$ under different SNP-selection strategies. For each LD threshold, random thinning is compared with single-trait lead-SNP selection and MetaUSAT-based lead-SNP selection. Trait-wise counts were computed by summing, over all APCH discoveries, whether the selected subset contained each trait.

| LD threshold | Model | $L = 3$ | $L = 4$ | $L = 5$ | Runtime (min) |
|---|---|---|---|---|---|
| $r^2 = 0.10$ | APCH-b | $14.8 \pm 3.2$ | $3.9 \pm 1.9$ | $0.6 \pm 0.6$ | $0.27 \pm 0.02$ |
| | APCH-s | $35.7 \pm 4.7$ | $15.0 \pm 3.4$ | $5.2 \pm 2.1$ | $0.6 \pm 0.1$ |
| | APCH-$s_{full}$ | $35.8 \pm 4.2$ | $14.5 \pm 3.3$ | $4.8 \pm 1.9$ | $14.6 \pm 3.5$ |
| $r^2 = 0.25$ | APCH-b | $18.3 \pm 2.3$ | $5.4 \pm 1.5$ | $0.7 \pm 0.6$ | $0.44 \pm 0.05$ |
| | APCH-s | $48.6 \pm 5.4$ | $19.5 \pm 3.5$ | $8.4 \pm 1.9$ | $1.1 \pm 0.2$ |
| | APCH-$s_{full}$ | $48.6 \pm 5.2$ | $19.1 \pm 3.2$ | $8.3 \pm 1.6$ | $39.5 \pm 10.5$ |
| $r^2 = 0.60$ | APCH-b | $20.6 \pm 2.0$ | $8.3 \pm 1.6$ | $1.4 \pm 0.5$ | $0.91 \pm 0.09$ |
| | APCH-s | $67.9 \pm 5.0$ | $25.6 \pm 2.4$ | $11.6 \pm 1.4$ | $2.6 \pm 1.8$ |
| | APCH-$s_{full}$ | $67.3 \pm 4.6$ | $24.7 \pm 2.4$ | $11.7 \pm 1.4$ | $89.9 \pm 28.9$ |

Supplementary Table 5: Comparison of APCH-b, APCH-s, and APCH-$s_{full}$ under random LD thinning. APCH-$s_{full}$ refers to the APCH-s model without pruning allocation states. Entries are mean $\pm$ standard deviation across 50 replicates for tag SNP discoveries at subset levels $L = 3$, $L = 4$, and $L = 5$, using subset-level lfsr at $\alpha = 0.01$. The average numbers of retained SNPs were 109,650 for $r^2 = 0.10$, 181,641 for $r^2 = 0.25$, and 354,540 for $r^2 = 0.60$. All analyses were run on a Linux machine (AlmaLinux 8.10) with R (version 4.5.2) linked to OpenBLAS (version 0.3.15). We used 8 CPU threads per run. Under the threshold ($r^2 = 0.25$), a typical APCH-s run for 5 GWAS traits required about 2 GB of memory.

| Model | Inference | Tag SNP count | $P(\min\lvert z\rvert < 2)$ | PROI | FI | T2D | FPG | 2hPG |
|---|---|---|---|---|---|---|---|---|
| APCH-s | FSR | 8.4 ± 1.9 | 0.000 | 3.79 | 3.95 | 9.45 | 9.12 | 3.80 |
| APCH-s | FDR | 59.4 ± 29.4 | 0.781 | 2.40 | 2.21 | 6.93 | 4.02 | 2.86 |
| APCH-b | FSR | 0.7 ± 0.6 | 0.000 | 12.17 | 4.70 | 39.38 | 13.06 | 9.08 |
| APCH-b | FDR | 20.0 ± 8.0 | 0.787 | 6.39 | 1.54 | 3.40 | 4.03 | 1.64 |

Supplementary Table 6: Comparison of $L = 5$ discoveries under lfsr- and lfdr-based inference at $r^2 = 0.25$. Entries compare APCH-s and APCH-b at $\alpha = 0.01$. Tag SNP count is reported as mean ± standard deviation across 50 random-thinning replicates. The weak-coordinate proportion $P(\min\lvert z\rvert < 2)$ is computed as the pooled proportion of $L = 5$ tag SNPs with at least one trait having $\lvert z\rvert < 2$. Trait-specific columns report the median absolute $z$-score among all $L = 5$ tag SNPs.

| Trait | Grids after pruning | Grids after distillation | $\pi_0$ | Conditional non-null distribution |
|---|---|---|---|---|
| PROI | 5 | 2 | 0.9938 | $0.933\,N(0, 2.06^2) + 0.067\,N(0, 9.02^2)$ |
| FI | 5 | 2 | 0.8747 | $0.978\,N(0, 0.81^2) + 0.022\,N(0, 2.72^2)$ |
| T2D | 6 | 3 | 0.6126 | $0.954\,N(0, 0.88^2) + 0.044\,N(0, 2.71^2) + 0.002\,N(0, 7.72^2)$ |
| FPG | 8 | 3 | 0.6868 | $0.982\,N(0, 0.57^2) + 0.017\,N(0, 3.11^2) + 0.001\,N(0, 10.84^2)$ |
| 2hPG | 4 | 2 | 0.8248 | $0.989\,N(0, 0.45^2) + 0.011\,N(0, 2.21^2)$ |
| Tissue | | | | |
| SAT | 3 | 1 | 0.9593 | $1.000\,N(0, 2.06^2)$ |
| VAT | 2 | 2 | 0.9585 | $0.973\,N(0, 2.01^2) + 0.027\,N(0, 4.95^2)$ |
| Muscle | 3 | 2 | 0.9445 | $0.964\,N(0, 1.83^2) + 0.036\,N(0, 4.95^2)$ |
| Pancreas | 2 | 2 | 0.9655 | $0.951\,N(0, 2.01^2) + 0.049\,N(0, 4.95^2)$ |
| Blood | 2 | 1 | 0.9568 | $1.000\,N(0, 2.08^2)$ |
| Liver | 2 | 1 | 0.9684 | $1.000\,N(0, 2.07^2)$ |

Supplementary Table 7: Marginal ash fits and grid distillation for GWAS traits in the primary analysis and for TWAS tissues. For each trait or tissue, we report the number of non-null slabs retained after pruning, the number retained after grid distillation, the estimated null proportion $\pi_0$, and the conditional non-null distribution. Slabs were pruned when their conditional non-null weight was less than $10^{-6}$.

| Gene | $L$ | lfsr | SAT | VAT | Muscle | Pancreas | Blood | Liver |
|---|---|---|---|---|---|---|---|---|
| *JAZF1* | 5 | $1.43 \times 10^{-5}$ | **-7.52** | **-9.27** | **-10.13** | **-9.55** | 1.33 | **-5.71** |
| *WFS1* | 3 | $1.07 \times 10^{-6}$ | **8.48** | **8.09** | **7.88** | -2.23 | – | – |
| *ANK1* | 3 | $2.75 \times 10^{-3}$ | **5.90** | – | **8.42** | **-3.30** | – | – |
| *IRS1* | 2 | $9.87 \times 10^{-6}$ | **-7.30** | **-6.33** | – | 2.37 | 1.35 | – |
| *ST6GAL1* | 2 | $2.17 \times 10^{-3}$ | – | – | -2.10 | **6.02** | **3.70** | – |
| *TIGAR* | 2 | $2.29 \times 10^{-3}$ | **-3.81** | – | -1.29 | -1.91 | **-4.72** | – |
| *ARPIN* | 2 | $4.78 \times 10^{-3}$ | **4.18** | **5.43** | -0.78 | – | 0.92 | – |
| *SCX* | 2 | $9.20 \times 10^{-3}$ | **4.96** | **-2.68** | – | – | – | – |
| *LINC00623* | 2 | $1.71 \times 10^{-2}$ | -1.19 | -1.24 | – | **2.55** | **-5.41** | -0.23 |
| *TPH1* | 2 | $2.11 \times 10^{-2}$ | **2.67** | 1.79 | **-2.26** | -1.88 | – | -0.18 |

Supplementary Table 8: Jointly significant genes ($L \geq 2$) in the multi-tissue TWAS of T2D, identified by APCH using lfsr at $\alpha = 0.01$. Calibrated $z$-scores are shown for the six tissues; tissues in the selected subset are in bold, and – indicates that the gene is not predictable in that tissue.

### 3 Supplementary Figures

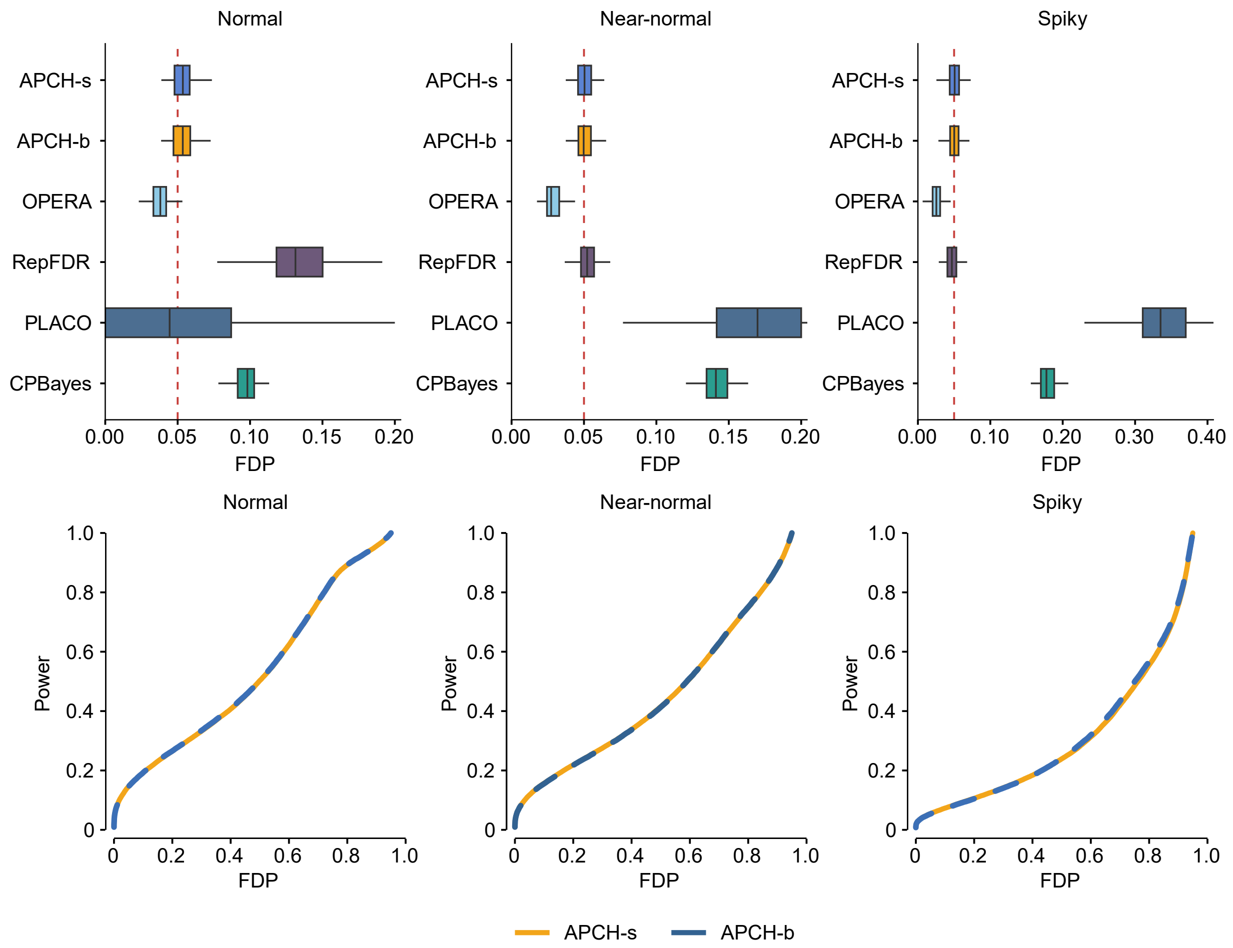


Supplementary Figure 1: **False discovery control and power in Simulation Study 1.** This figure uses the mixed-configuration setting in Simulation Study 1, $(\varphi_{00}, \varphi_{10}, \varphi_{01}, \varphi_{11}) = (0.80, 0.10, 0.05, 0.05)$. The top row shows the distribution of the realized FDP at $\alpha = 0.05$. The bottom row shows power versus FDP curves for APCH-s and APCH-b. Results are averaged over 100 replicates. PLACO estimates nuisance parameters by assigning marginal null/non-null labels with $c = 10^{-4}$, treating $p_{ij} < c$ as non-null and $p_{ij} > c$ as null for trait $j$. This gives groups $S_{ab}$, $a, b \in \{0, 1\}$, such as $S_{10} = \{i : p_{i1} < c,\ p_{i2} > c\}$. The mean $\pm$ SD counts in $(S_{01}, S_{10}, S_{11})$ were $178 \pm 15, 278 \pm 19, 20 \pm 4$, $175 \pm 14, 272 \pm 18, 18 \pm 3$, and $140 \pm 12, 218 \pm 14, 11 \pm 3$ for the Normal, Near-normal, and Spiky scenarios, respectively. In each boxplot, the center line denotes the median, the box bounds denote the 25th and 75th percentiles, and the whiskers extend to 1.5 times the interquartile range.

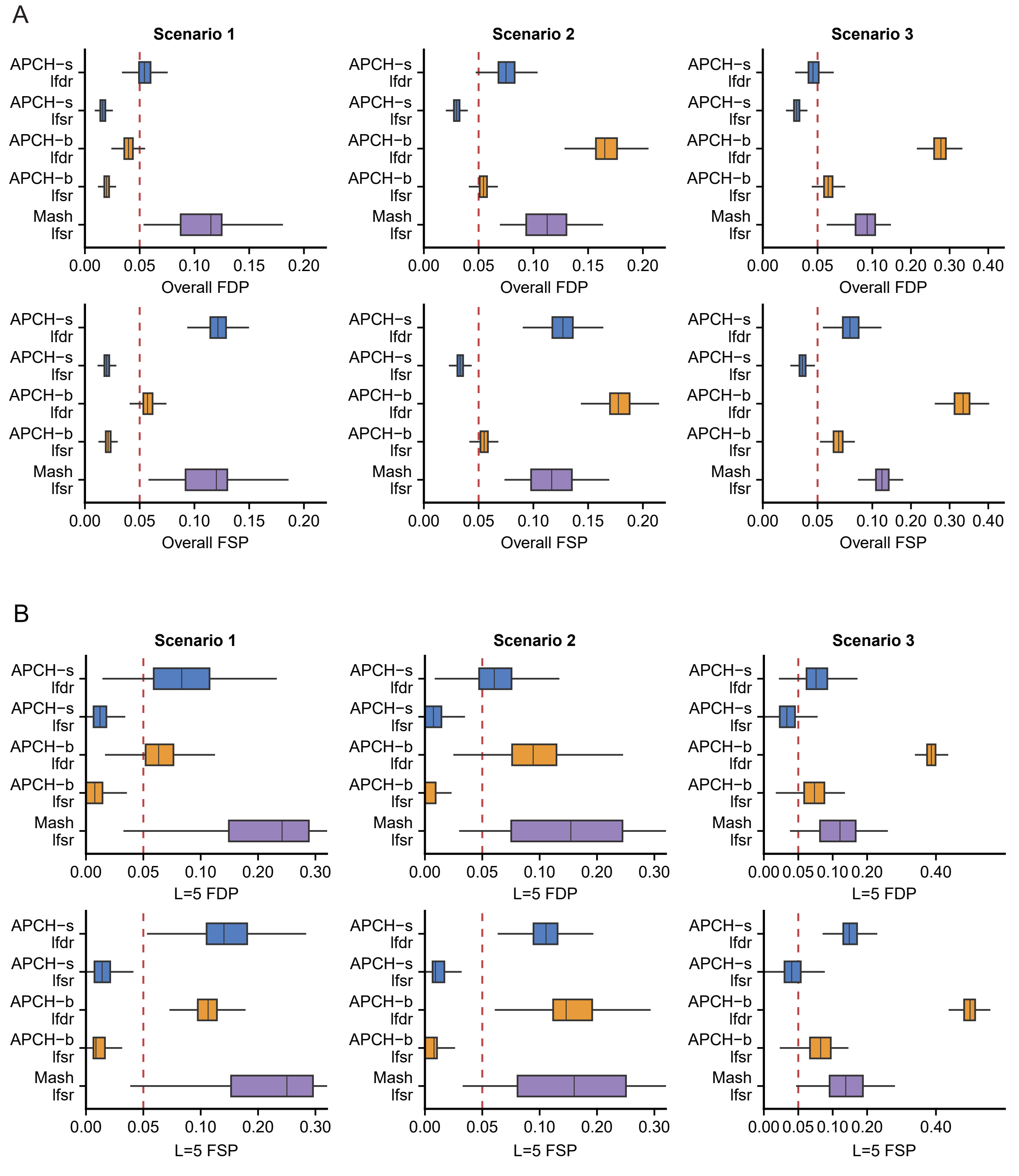


Supplementary Figure 2: **Distribution of FDP and FSP in Simulation Study 2.** Boxplots summarize the false discovery proportion (FDP) and false sign proportion (FSP) across 500 simulation replicates. FDP measures the proportion of reported subsets that include at least one truly null feature. FSP additionally counts boundary discoveries with incorrect inferred signs as errors. In each boxplot, the center line denotes the median, the box bounds denote the 25th and 75th percentiles, and the whiskers extend to 1.5 times the interquartile range.

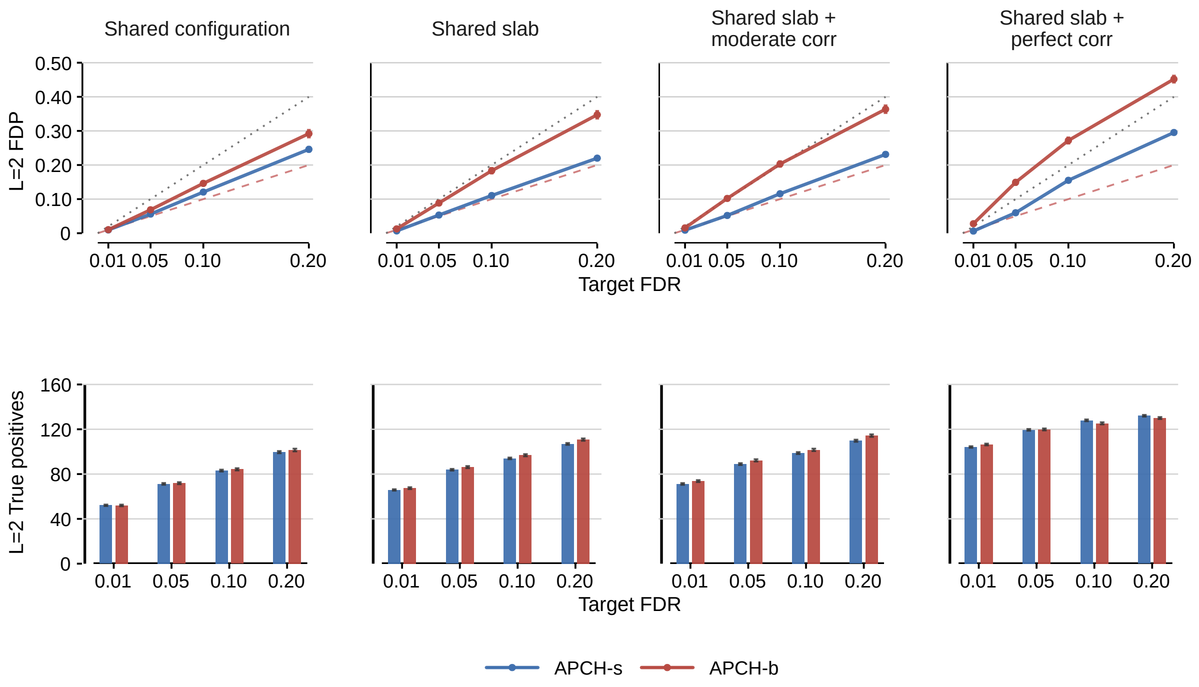


Supplementary Figure 3: **Comparison of APCH-b and APCH-s in Simulation Study 3.** $L = 2$ FDP and true positive discoveries under different sharing structures using lfsr-based inference. Error bars show standard errors over 100 replicates. The red dashed line denotes $y = x$ and the gray dashed line denotes $y = 2x$. APCH-s is more robustly calibrated than APCH-b across sharing structures while retaining comparable power. It is interesting that APCH-b does not perform well in the first scenario. This is because the effects are comparatively more sparse than in Simulation 1.

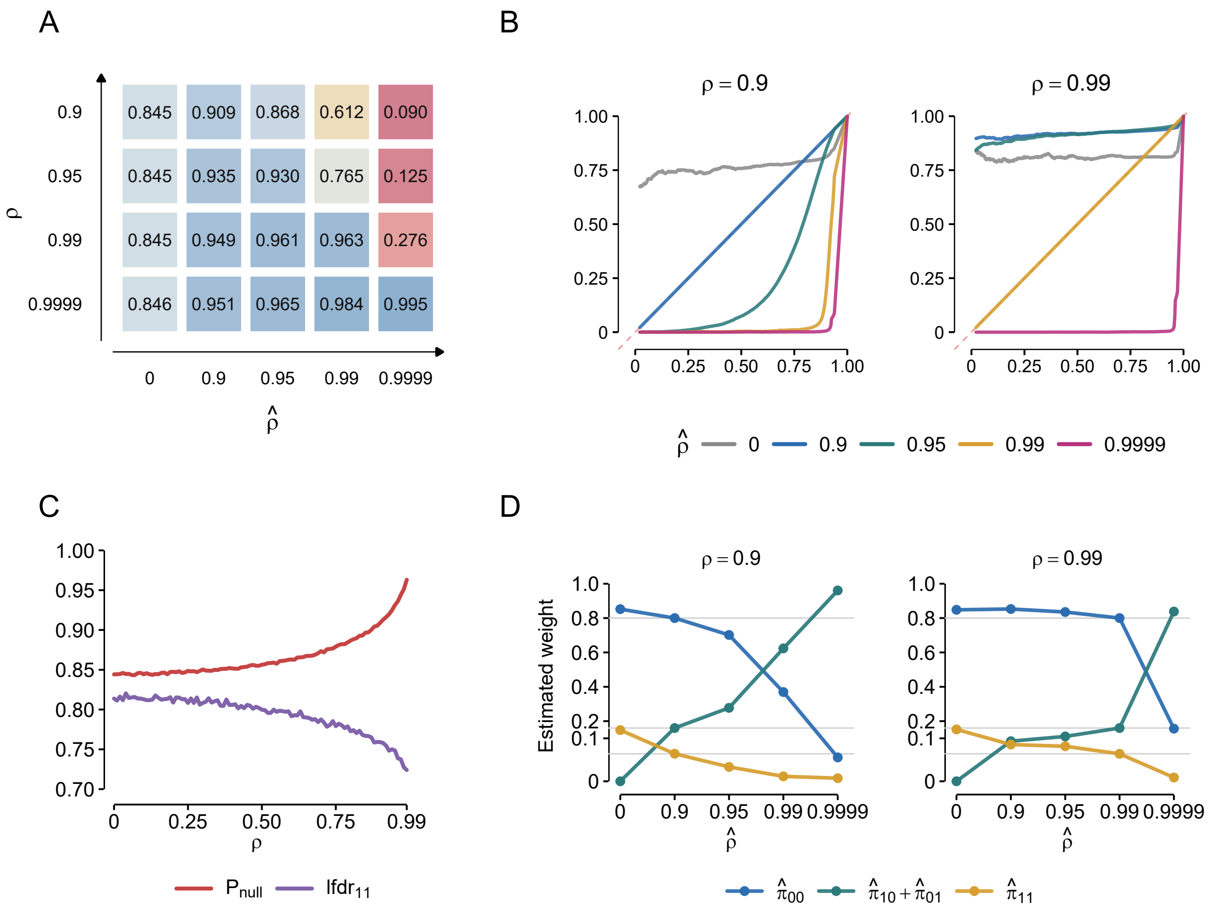


Supplementary Figure 4: **Effect of misspecifying the estimation-error correlation on APCH posterior quantities and prior-weight estimation.** (A) Mean posterior probability of the global-null configuration, $P(\gamma = 00 \mid \hat{b}_i)$, under global-null data generated with different true estimation-error correlations $\rho$ and evaluated using plug-in correlations $\hat{\rho}$. (B) Calibration curves comparing $P(\gamma = 00 \mid \hat{b}_i)$ computed under the correctly specified correlation (X-axis) with that computed under misspecified $\hat{\rho}$ (Y-axis). (C) Mean posterior $P(\gamma = 00 \mid \hat{b}_i)$ for observations generated from the global null and mean $\text{lfdr}_{1,2}$ for observations generated from the joint non-null configuration as a function of the correctly specified $\rho$. (D) Estimated prior weights from refitting only the configuration proportions under misspecified $\hat{\rho}$, with the slab variance fixed. Across simulations, the true prior was $(\pi_{00}, \pi_{10}, \pi_{01}, \pi_{11}) = (0.80, 0.08, 0.08, 0.04)$.

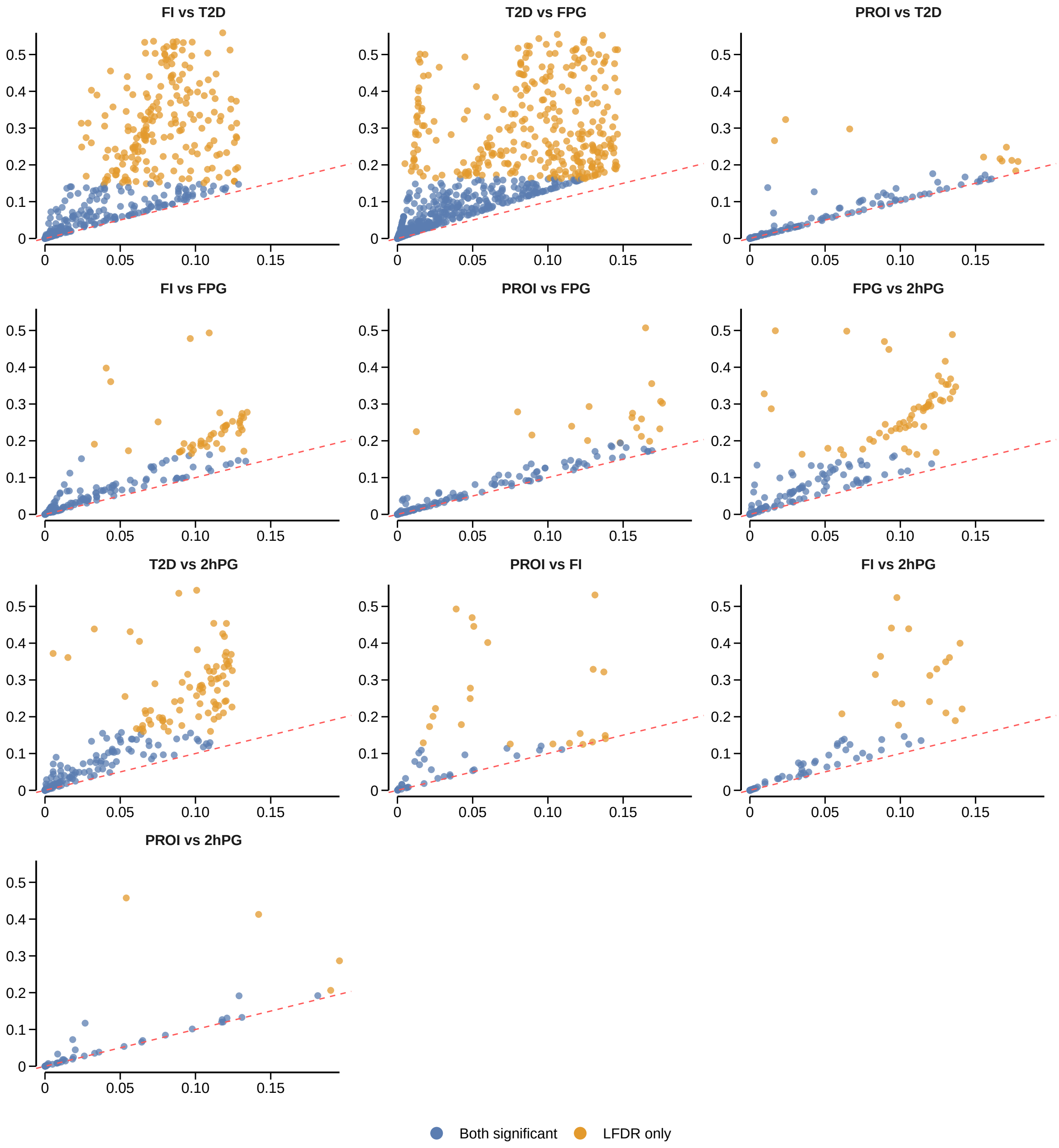


Supplementary Figure 5: **Comparison of lfdr and lfsr for pairwise GWAS traits.** APCH was run separately for each trait pair. Each point represents a SNP declared jointly significant for the corresponding trait pair using lfdr at $\alpha = 0.05$. Blue points are significant under both lfdr- and lfsr-based inference, whereas orange points are significant under lfdr only. The red dashed line denotes $y = x$.

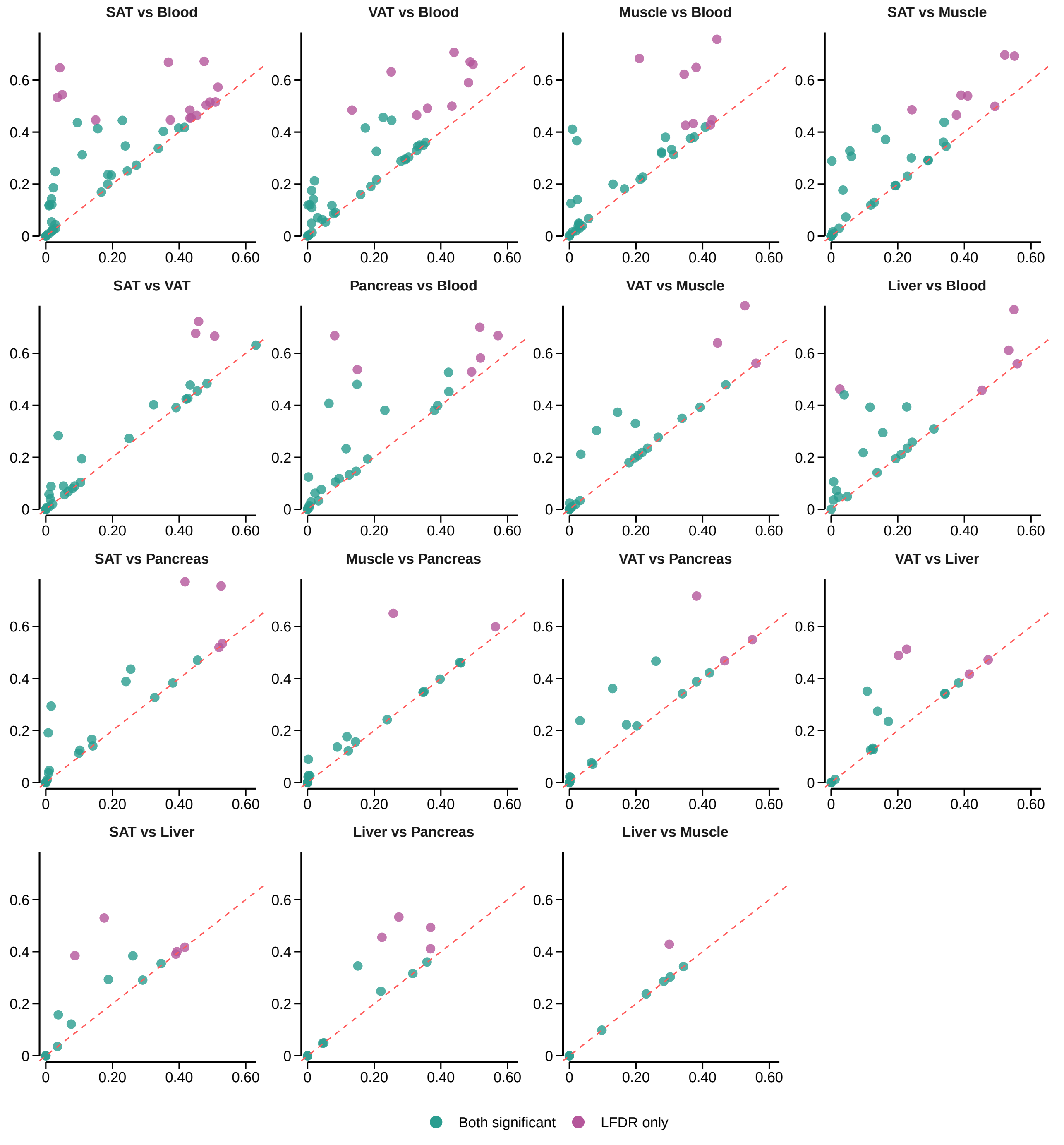


Supplementary Figure 6: **Comparison of lfdr and lfsr for pairwise TWAS tissues.** APCH was run separately for each pair of the six tissues. Each point represents a gene declared jointly significant for the corresponding tissue pair using lfdr at $\alpha = 0.2$. Green points are significant under both lfdr- and lfsr-based inference, whereas purple points are significant under lfdr only. The red dashed line denotes $y = x$.

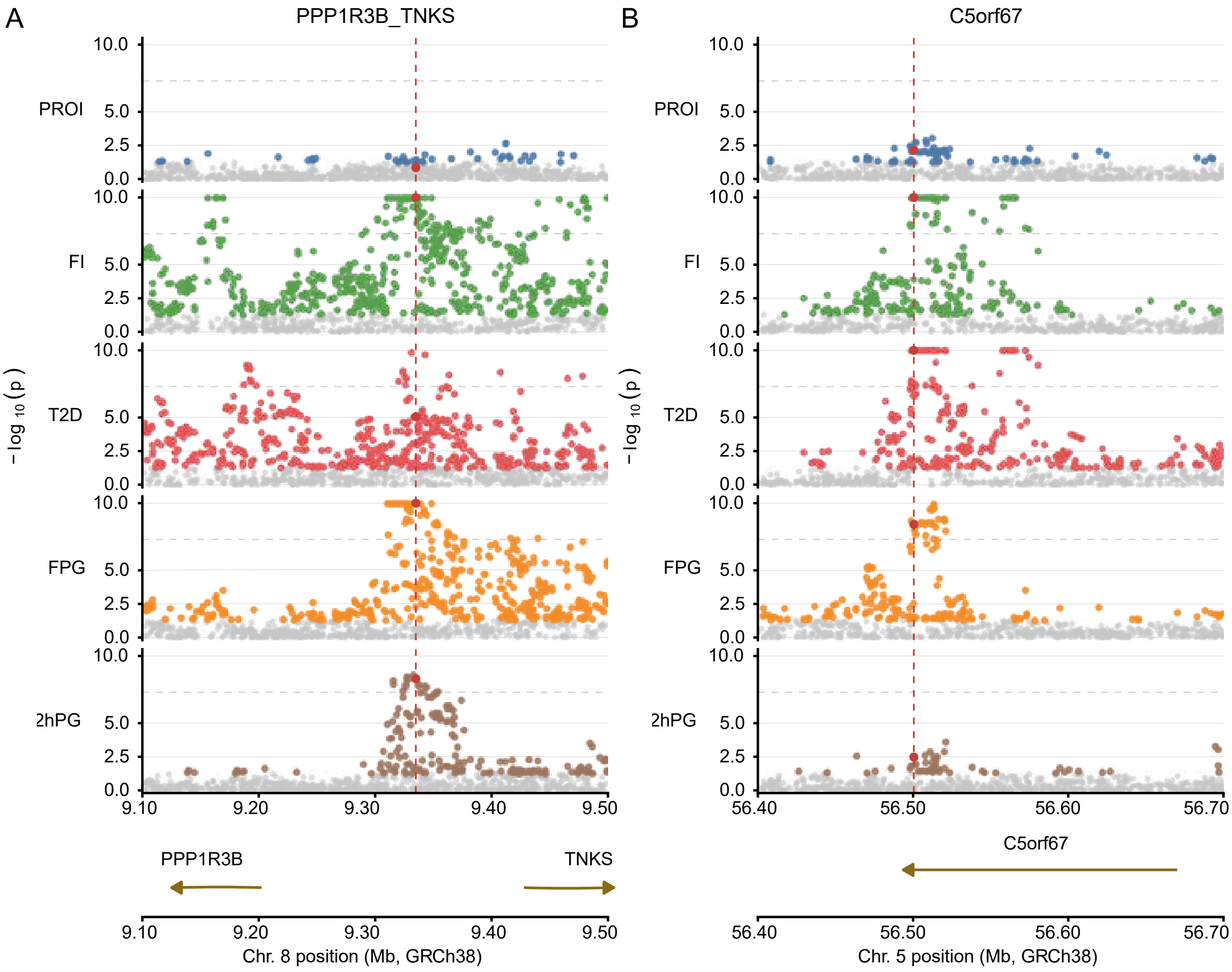


Supplementary Figure 7: **Regional association plots for APCH-selected loci.** Panel A shows a partial-trait association near *PPP1R3B*/*TNKS*, for which APCH inferred joint significance across four traits, excluding PROI. Panel B shows a less well-known but literature-supported [26, 27] locus near *C5orf67*, with joint significance across all five traits. Shown are association $p$-values from five traits, with $-\log_{10}(p)$ truncated at 10. Red points indicate the SNPs selected by APCH for prior fitting.
